# Striatal lateral inhibition regulates abnormal movements in a mouse model of levodopa-induced dyskinesia

**DOI:** 10.1101/2024.10.11.617939

**Authors:** Emily L. Twedell, Olivia K. Barnhill, Chloe J. Bair-Marshall, Allison E. Girasole, Lara K. Scaria, Sadhana Sridhar, Alexandra B. Nelson

**Affiliations:** Neuroscience Graduate Program, UCSF, San Francisco, CA 94158, USA; Kavli Institute for Fundamental Neuroscience, UCSF, San Francisco, CA 94158, USA; Weill Institute for Neurosciences, UCSF, San Francisco, CA 94158, USA; Department of Neurology, UCSF, San Francisco, CA 94158, USA; Aligning Science Across Parkinson’s (ASAP) Collaborative Research Network, Chevy Chase, MD 20815, USA

## Abstract

Striatal medium spiny neurons (MSNs) integrate convergent cortical, thalamic, and dopaminergic inputs to shape motor output. In addition, MSNs form local inhibitory synaptic connections with one another. Striatal lateral inhibition may contribute to the regulation of motor output, but its role in pathological motor function remains incompletely understood. Disruptions in motor function characterize several movement disorders, including levodopa-induced dyskinesia (LID), a complication of Parkinson’s disease (PD) therapy characterized by involuntary movements. Here, we identify chronic changes in the strength of striatal lateral inhibitory synapses in a mouse model of PD/LID. These synapses are also modulated by acute dopamine signaling. Chemogenetic suppression of lateral inhibition originating from dopamine D2 receptor-expressing MSNs lowers the threshold for the expression of involuntary movements *in vivo*. Together, these findings identify a contribution of striatal lateral inhibition to motor dysfunction and suggest a broader role for this microcircuit in regulating motor output.

## INTRODUCTION

Motor control, including the initiation and coordination of movement, relies on the activity of basal ganglia circuitry. As the input nucleus of the basal ganglia, the striatum integrates excitatory input from the cortex and neuromodulatory input from midbrain dopamine neurons. The striatum consists primarily of GABAergic projection neurons, also known as medium spiny neurons (MSNs)(*1*, *2*). MSNs can be divided into two subpopulations: dopamine D1 receptor-expressing (D1-MSN) and D2 receptor-expressing (D2-MSN) neurons. D1- and D2-MSNs are coactivated during movement initiation(*3–6*), but some studies have identified distinct functions in action selection and motor learning (*5–8*). One parsimonious model is that ensembles of D1-MSNs facilitate a desired behavior while related ensembles of D2-MSNs suppress competing behaviors (*9*, *10*).

One way to better understand how striatal circuits coordinate motor output is to examine conditions in which motor function is disrupted. One such condition is levodopa-induced dyskinesia (LID), a complication of Parkinson’s disease (PD) in which treatment with the dopamine precursor levodopa results in abnormal, involuntary movements. Prior studies suggest aberrant coordination of D1- and D2-MSN activity in PD/LID, including suppressed D2-MSN activity, abnormally high D1-MSN activity, maladaptive corticostriatal connectivity, and larger movement-related D1-MSN ensembles with levodopa treatment (*6*, *11–14*).

How are active ensembles of D1-MSNs and D2-MSNs shaped at the synaptic level? Within the striatum, MSNs receive several inhibitory inputs, including from local GABAergic interneurons, and from other MSNs (*2*, *15–18*). The potential impact of MSN-MSN lateral inhibition has been questioned due to low connection rates and small unitary inhibitory currents (*19*). However, since MSNs represent 95% of all striatal neurons, these connections may collectively have a substantial impact on circuit function. Previous studies have demonstrated that interactions between D1- and D2-MSN populations contribute to locomotor behavior and goal-directed learning (*20*, *21*). However, how local MSN-MSN inhibitory synaptic transmission contributes to motor dysfunction, particularly in the context of parkinsonism and levodopa-induced dyskinesia, remains unclear. In sensory systems, lateral inhibition plays a role in improving sensory perception by shaping the pattern of neural activity evoked by sensory stimuli (*22–29*). This raises the question of whether similar inhibitory motifs could contribute to the coordination of MSN ensemble activity during motor behavior (*9*).

To investigate the potential contribution of striatal lateral inhibition to motor dysfunction, we used *ex vivo* electrophysiology and chemogenetics in a mouse model of PD/LID. We found remodeling of D2- and D1-MSN lateral connections onto D1-MSNs in the mouse model of PD/LID. Chemogenetic inhibition of D2-mediated striatal lateral inhibition lowered the threshold for dyskinesia. Together, these findings demonstrate that disruption of striatal lateral inhibition can exacerbate pathological motor output in the context of dopamine depletion and chronic levodopa treatment. These results identify a circuit-level mechanism that modulates dyskinesia severity and suggest that local inhibitory interactions may play a broader role in regulating striatal output.

## RESULTS

### Striatal lateral connections between MSNs are asymmetric

To study MSN-MSN inhibitory synaptic connections, we employed an optogenetic strategy (*30*). We injected a Cre-dependent channelrhodopsin into the dorsolateral striatum (DLS) of A2a-Cre;Drd1-tdTomato or Drd1-Cre;Drd1-tdTomato mice (Figure 1A). We focused on the DLS because of its established role in sensorimotor function (*31*) and causal role in the expression of levodopa-induced dyskinesia (*32*). In D1-Cre mice, D1-MSN to D1-MSN (D1-D1) and D1-MSN to D2-MSN (D1-D2) connections can be assayed, while in A2a-Cre mice, D2-MSN to D1-MSN (D2-D1) and D2-MSN to D2-MSN (D2-D2) connections can be assayed. We prepared acute brain slices containing the striatum and performed whole-cell voltage-clamp recordings of D1-MSNs (tdTomato-positive) or D2-MSNs (tdTomato-negative) in the DLS (Figure 1B). Optical stimulation evoked inhibitory postsynaptic currents (oIPSCs), which represent the sum of local lateral inhibitory inputs onto the post-synaptic neuron (Figure 1C, inset). These oIPSCs were GABA_A_ receptor-dependent, as they were blocked by picrotoxin (S1A-B). oIPSCs were evoked at several light powers and compared across the four MSN-MSN connection types (Figure 1C-F). A prior study using paired MSN-MSN recordings found that synaptic connections were asymmetric; those arising from D2 neurons had higher connection strengths (*19*). Consistent with these findings, we found that D2-D1 oIPSCs had the highest amplitudes (Figure 1G). There were no significant differences in oIPSC amplitude across the other MSN-MSN connection types. These experiments indicate that the optical approach can recapitulate the asymmetry in MSN-MSN synaptic connectivity, with D2-D1 lateral inhibition being the strongest.

**Figure 1.**
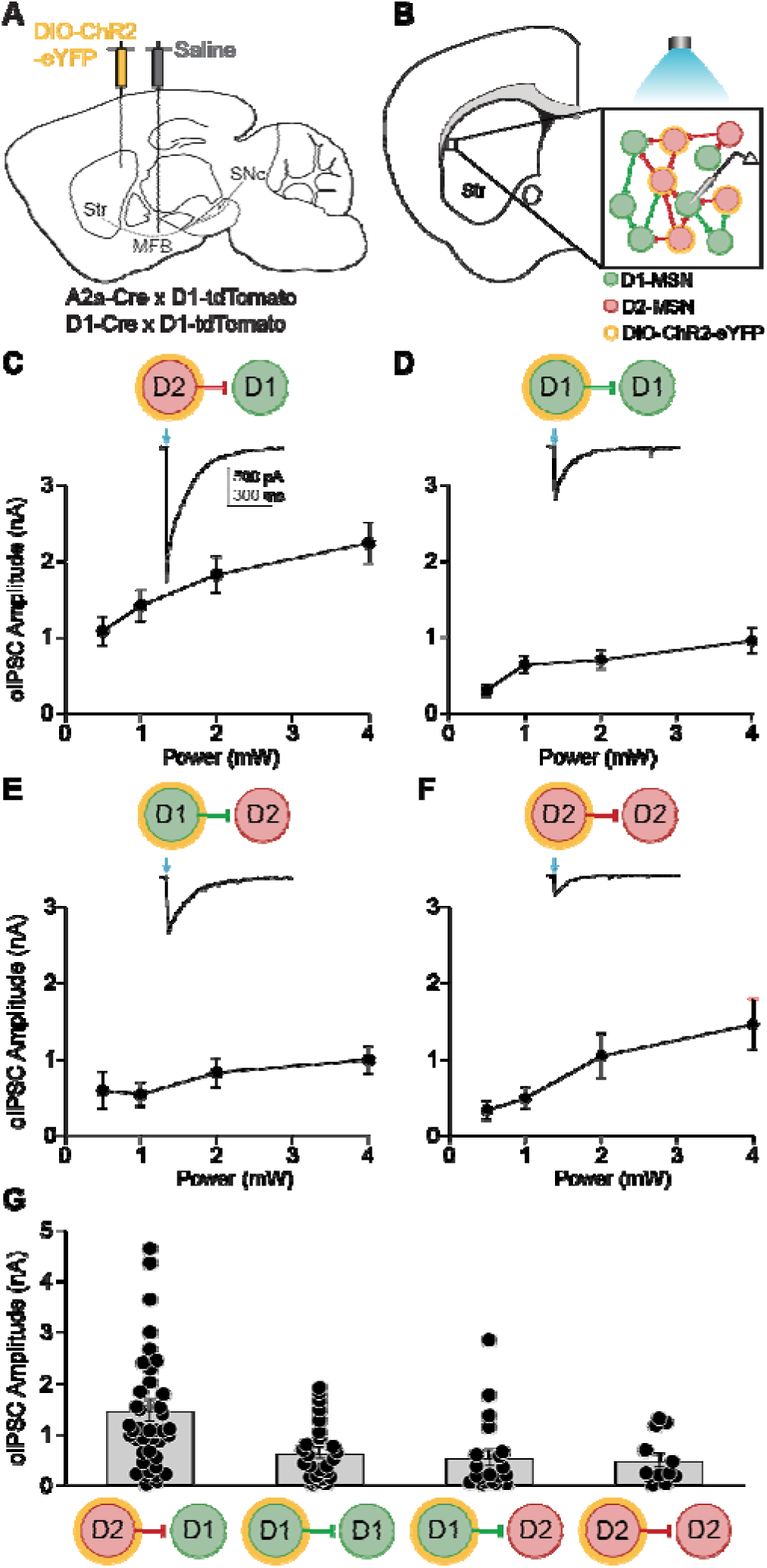
Striatal MSN-MSN synaptic connectivity is asymmetric. **(A)** Sagittal schematic showing DIO-ChR2-eYFP injection into the dorsolateral striatum (DLS) and saline injection into the medial forebrain bundle (MFB). **(B)** Coronal schematic depicting an *ex vivo* slice containing the DLS. *Inset*: In this example depicting an A2a-Cre; D1-tdTomato mouse, a whole-cell recording of a medium spiny neuron (D1-MSN, green) is made. Synaptic currents were optically evoked from local D2-MSN (red) collaterals expressing ChR2 (yellow). **(C)** – **(F)** Average oIPSC amplitude evoked by 0.5, 1, 2, and 4 mW light pulses. *Insets*: Representative oIPSCs recorded from MSNs during optical activation of local collaterals. (C) D2-D1 (N = 10, n = 38). (D) D1-D1 (N = 5, n = 26). (E) D1-D2 (N = 8, n = 19). (F) D2-D2 (N = 6, n = 13). **(G)** Average oIPSC amplitude in response to 1 mW optical stimulation for all MSN-MSN connection types. D2-D1: N = 12, n = 41; D1-D1: N = 5, n = 26; D1-D2: N = 9, n = 20; D2-D2: N = 6, n = 12. KW: p = 0.0002. Each dot represents data from one neuron. N = animals; n = cells. Data shown as mean ± SEM.

### D2-D1 striatal lateral connections are remodeled in parkinsonian animals and parkinsonian animals chronically treated with levodopa

Loss of dopamine in the parkinsonian state and treatment with chronic levodopa trigger significant changes in basal ganglia activity (*33*). These alterations in activity are thought to drive homeostatic changes in excitability and synaptic connectivity (*34–36*). We hypothesized that this homeostatic regulation may also take place within the striatal lateral inhibitory network.

To understand how lateral inhibition may contribute to motor dysfunction, we tested MSN-MSN connectivity in a mouse model of Parkinson’s disease (PD) and levodopa-induced dyskinesia (LID). These conditions are associated with distinct abnormalities in motor output: PD is characterized by impaired movement initiation and bradykinesia, whereas LID is characterized by abnormal, involuntary movements. To model these disorders, we used the 6-OHDA mouse model of PD. The neurotoxin 6-OHDA was injected unilaterally in the medial forebrain bundle (Figure 2A), resulting in severe dopamine depletion (Figure 2B). Control animals were injected intracranially with saline. Animals recovered for several weeks, after which they were randomized to daily intraperitoneal (IP) injections of levodopa or saline (Figure 2C). 6-OHDA treated animals (Park and LD OFF) exhibited a reduction in movement velocity and an ipsilesional rotational bias (Figure 2D; S2). Parkinsonian animals injected with levodopa (5 mg/kg; LD ON) showed increased movement velocity, contralateral rotation bias, and dyskinesia (Figure 2D-E; S2). In response to each levodopa injection, animals developed abnormal involuntary movements (AIMs) which lasted 60-100 minutes, after which animals resumed baseline movement (Figure 2E). These manipulations yielded 3 key experimental groups for subsequent slice electrophysiology (MFB injection/Daily IP injection): Control (saline/saline), Park (6OHDA/saline) and LD (6OHDA/LD).

**Figure 2.**
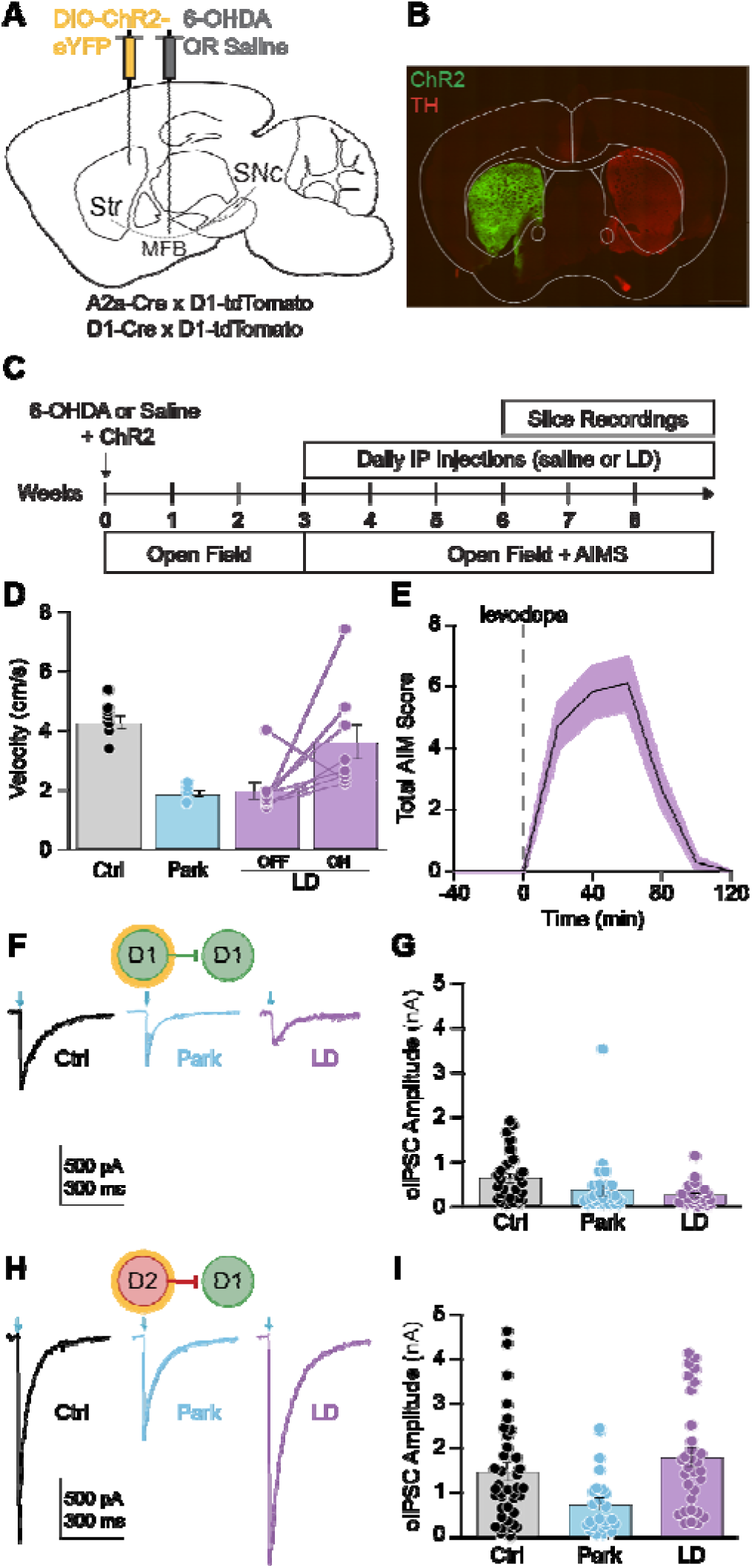
Dopamine depletion and chronic levodopa treatment remodel D2-D1 striatal lateral connections. **(A)** Sagittal schematic showing DIO-ChR2-eYFP injection into the dorsolateral striatum (DLS) and 6-OHDA or saline injection into the medial forebrain bundle (MFB). **(B)** Representative coronal section of the DLS showing DIO-ChR2-eYFP expression and loss of TH immunoreactivity in the ipsilesional hemisphere of a 6-OHDA-treated mouse. TH was imaged with a secondary antibody at 647 nm, but is false-colored red for visibility. The tdTomato channel is not shown. Scale bar is 1 mm. **(C)** Experimental timeline. **(D)** Average velocity in the open field, measured over a 10-minute recording period beginning 20 minutes after intraperitoneal (IP) injection of saline (Ctrl and Park) or levodopa (LD; 5 mg/kg). Ctrl: N = 9; Park: N = 5; LD: N = 8. **(E)** Mean abnormal involuntary movement (AIM) score following IP injection of levodopa (5 mg/kg). Total AIM scores were calculated by summing axial, limb, and orolingual dyskinesia scores (0-4 each; maximum total score of 12). Each animal was scored every 20 minutes, and values represent the mean across animals at each time point. LD: N = 11. **(F)** Representative D1-D1 oIPSCs recorded from animals in Ctrl (*left*), Park (*middle*), and LD (*right*) groups. **(G)** Average D1-D1 oIPSC amplitude in response to 1 mW optical stimulation for Ctrl (N = 5, n = 26), Park (N = 7, n = 28), and LD (N = 6, n = 24) groups. KW: p = 0.0142. **(H)** Representative D2-D1 oIPSC recorded from animals in Ctrl (*left*), Park (*middle*), and LD (*right*) groups. **(I)** Average D2-D1 oIPSC amplitude in response to 1 mW optical stimulation for Ctrl (N = 12, n = 41), Park (N = 6, n = 24), and LD (N = 8, n = 32) groups. KW: p = 0.0006. Each overlaid dot represents one cell (G, I). N = animals; n = cells. Data shown as mean ± SEM.

In the striatum, dopamine depletion leads to decreased D1-MSN activity (*6*, *12*). We hypothesized that this hypoactivity may lead to homeostatic downregulation of inhibitory connections onto D1-MSNs. Indeed, a study of paired recordings found D2-D1 and D1-D1 inhibitory responses to be markedly reduced in parkinsonian animals (*19*). To determine whether these connections are also weakened using the optical approach, we compared lateral inhibition in brain slices made from healthy and parkinsonian animals. Consistent with our hypothesis, oIPSC amplitudes were reduced for both D1-D1 (Figures 2F-G) and D2-D1 (Figures 2H-I) connections in Park animals relative to Ctrl.

In mouse models of PD, the activity of D2-MSNs is marginally, if at all, increased (*6*, *12*). Thus, homeostatic changes in lateral inhibition on to D2-MSNs may not be recruited. Due to low connection rates in healthy animals, Taverna et al. were unable to assess D1-D2 connections in the parkinsonian state (*19*). Here, we measured D1-D2 connectivity using the optical approach, finding no significant difference across groups (Figure S3A-B).

Less is known about homeostatic changes to synaptic strength in parkinsonian animals chronically treated with levodopa. During bouts of dyskinesia, D1-MSNs are hyperactive in rodent and non-human primate models of LID (*6*, *12*, *13*, *33*, *35*, *36*). We hypothesized that chronic treatment with levodopa may result in increased D2-D1 lateral inhibition to counteract excessive D1-MSN activity. To determine whether lateral inhibition onto D1-MSNs is altered by dopamine depletion and chronic levodopa treatment, we again used the optical approach. Brain slices were made 24h after the last IP treatment. As predicted, D2-D1 oIPSC amplitudes were restored to control levels in LD animals (Figures 2H-I). We did not detect a change in D1-D1 or D1-D2 synaptic strength in the LD condition relative to the treatment-naïve condition (Figures 2F-G, Figure S3A-B). These experiments indicate that lateral inhibition onto D1-MSNs is reduced by dopamine depletion, and that chronic levodopa treatment restores D2-D1 connections.

#### D2-D1 synaptic responses are inhibited by acute dopamine signaling

Chronic changes in neural activity resulting from dopamine depletion and replacement may drive homeostatic changes in synaptic connectivity. However, these changes may be maladaptive, creating vulnerability to acute changes in striatal dopamine. The previous experiments captured chronic changes in lateral inhibition, but do not recapitulate the acute effects of dopamine signaling. In LID, dyskinesia typically occurs at “peak-dose”, when brain levels of dopamine are elevated; as levodopa wears off, dyskinesia also resolves (Figure 2E) (*37*). Prior work indicates that dopamine acutely inhibits the amplitude of lateral inhibitory responses in *ex vivo* slices (*19*, *30*, *38–40*). These observations led to the hypothesis that acute levodopa treatment would lead to a reduction in the amplitude of D2-D1 synaptic responses.

To isolate the contribution of D2-receptor activation to the acute dopaminergic modulation of striatal lateral inhibition, we bath-applied the D_2_-agonist quinpirole while recording D2-D1 oIPSCs. We found that oIPSCs were acutely reduced by the agonist across all conditions (Figure 3A-C). As previously shown in healthy NAc, the reduction in oIPSC amplitude by the D_2_-agonist quinpirole is reversed by application of the D_2_-antagonist sulpiride (Figure S4), confirming its D2 receptor dependence (*30*, *38*). Taken together, these experiments demonstrate that in parkinsonian/levodopa-treated (LD) animals, acute dopamine signaling inhibits D2-D1 synaptic responses, which may in turn contribute to disinhibition of D1-MSNs *in vivo*.

**Figure 3.**
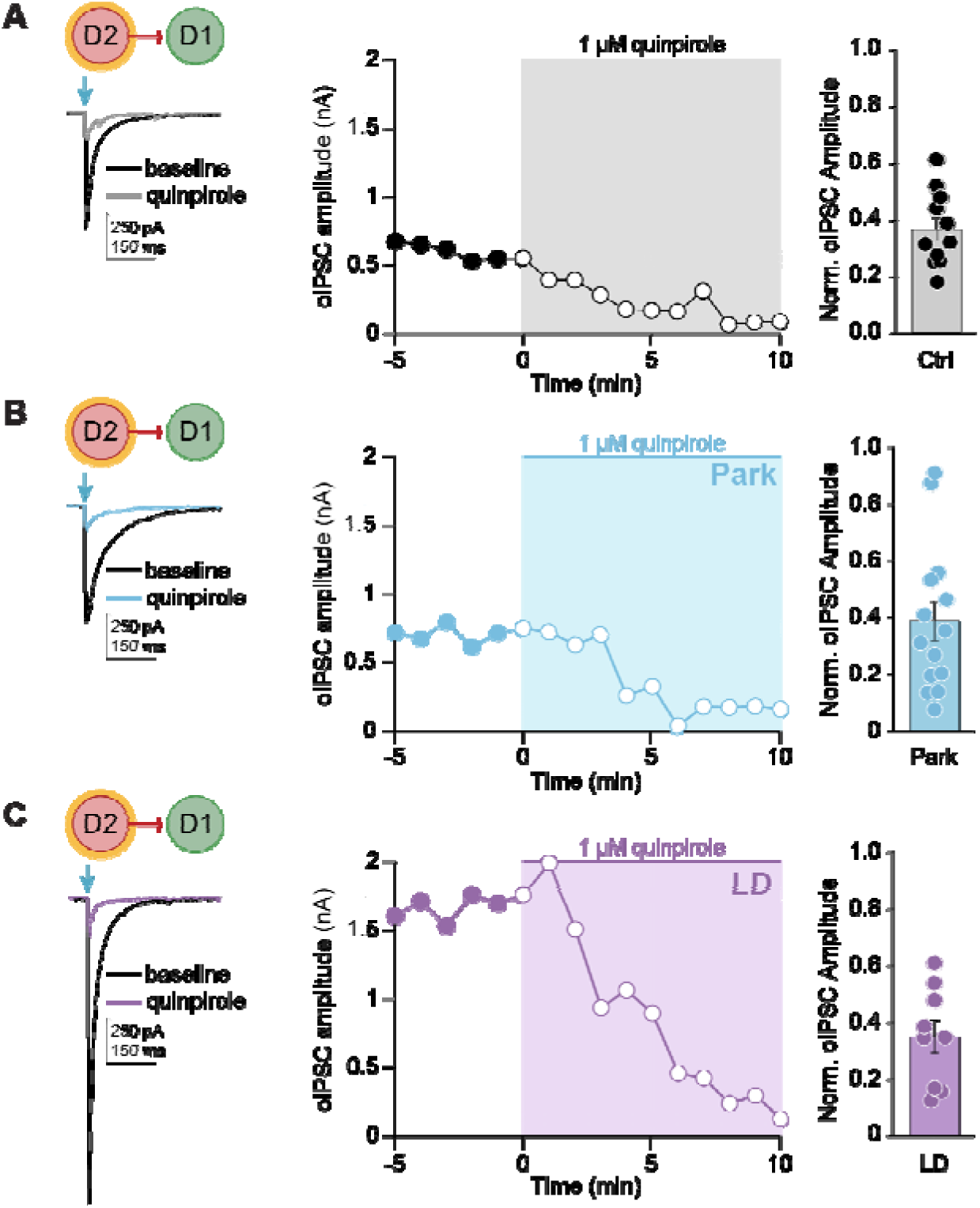
Dopamine signaling acutely inhibits D2-D1 synaptic responses. **(A) - (C)** (*Left*) Representative D2-D1 oIPSCs in Ctrl (A), Park (B), and LD (C) groups, before and after application of quinpirole (1 μM). (*Middle*) Time course of the effect of quinpirole on raw D2-D1 oIPSC amplitudes. (*Right*) Summary of D2-D1 oIPSC amplitudes in the presence of quinpirole, normalized to the mean pre-quinpirole baseline amplitude for each neuron (baseline = 1) for Ctrl (N = 6, n = 11, WSR: p = 0.0010), Park (N = 7, n = 14, p = 0.0001), and LD (N = 3, n = 9, WSR: p = 0.0039) groups. KW: p = 0.1004. Each dot represents data from one neuron. N = animals; n = cells. Data shown as mean ± SEM.

### Chemogenetic inhibition of D2-MSN-mediated striatal lateral inhibition lowers the threshold for levodopa-induced dyskinesia

While these results showed alterations in striatal lateral inhibition in *ex vivo* brain slices, we next aimed to test how lateral inhibition contributes to motor output *in vivo*. More specifically, we hypothesized that D2-D1 connections, the strongest source of MSN-MSN lateral inhibition onto D1-MSNs, may help constrain motor output by limiting excessive D1-MSN activity. Conversely, we hypothesized that a reduction in D2-D1 synaptic strength might lower the threshold for involuntary movements, as in LID. To date, it has been difficult to differentiate the effect of D2-MSNs on their intrastriatal (e.g., D1-MSNs) and extrastriatal (e.g., globus pallidus pars externa, GPe) targets, making it challenging to isolate the contribution of striatal lateral inhibition to motor output. To address this challenge, we employed a chemogenetic strategy. The inhibitory DREADD hM4D(G_i_) works via two primary mechanisms: (1) by pre-synaptic inhibition of release through G_i/o_-coupled signaling, and (2) by reducing excitability via G-protein-coupled inwardly rectifying potassium channels (GIRKs) (*41–44*). MSNs, however, do not express GIRKs (*45*, *46*), and thus local striatal activation of hM4D(G_i_) would be predicted to reduce local synaptic release (lateral inhibition) without changing the excitability/firing rate of the MSNs.

We validated this approach using both *ex vivo* and *in vivo* electrophysiology. We injected the DLS of A2a-Cre x D2-GFP mice with AAVs encoding Cre-dependent ChR2-eYFP and hM4D(G_i_)-mCherry (Figure 4A). Putative postsynaptic D1-MSNs were identified as non-fluorescent neurons, lacking green fluorescence (D2-GFP/ChR2-eYFP) and red fluorescence (hM4D(G_i_)-mCherry), while presynaptic D2-MSNs were optogenetically stimulated through their expression of ChR2. In acute striatal slices, we found that CNO markedly reduced the amplitude of D2-D1 oIPSCs, which represent compound inhibitory responses from optogenetically stimulated D2-MSNs (Figure 4B-C). Because only ChR2-expressing presynaptic neurons contribute directly to these optogenetically evoked responses, the reduction in oIPSC amplitude provides functional evidence that hM4D(G_i_) was expressed in at least a subset of the stimulated D2-MSNs, and when activated can suppress local D2-mediated synaptic inhibition. In separate current-clamp experiments, using A2a-Cre mice, we recorded from hM4D(G_i_)-mCherry positive D2-MSNs and found that CNO did not change their intrinsic excitability (Figure 4D-E; S5A-D).

**Figure 4.**
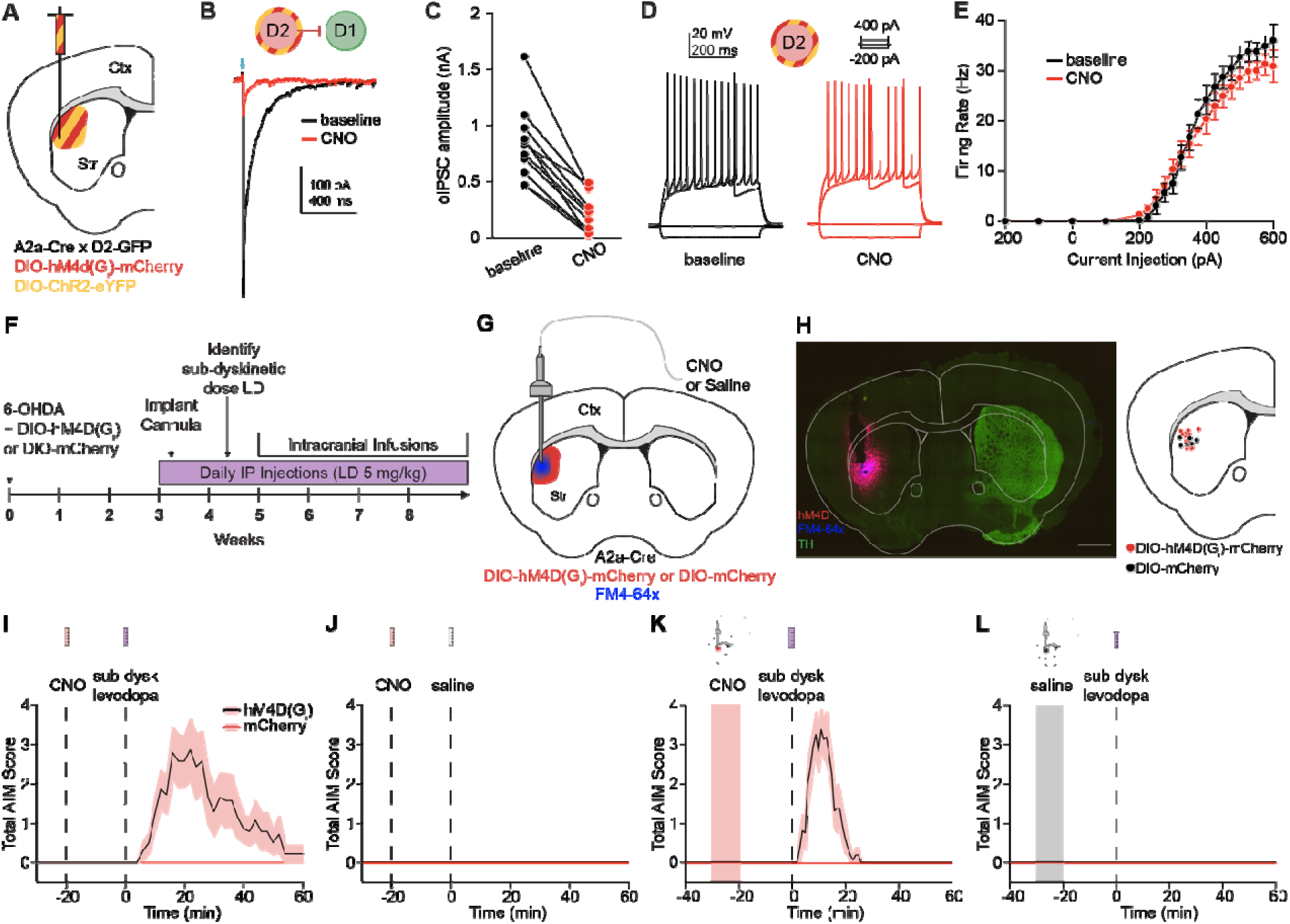
Chemogenetic inhibition of striatal lateral connections originating from D2-MSNs lowers the threshold for levodopa-induced dyskinesia. **(A)** Coronal schematic depicting co-injection of DIO-hM4D(G_i_)-mCherry and DIO-ChR2-eYFP in the DLS of an A2a-Cre x D2-GFP mouse. **(B)** Representative example of D2-D1 oIPSC before and after application of CNO. **(C)** Summary of changes in D2-D1 oIPSC amplitude with CNO application. N = 4, n = 11. WSR: p = 0.010. **(D)** Example firing rate of a D2-MSN in response to 500 ms steps of current injection before and after CNO application. **(E)** Summary current/firing (I/F) curves of D2-MSNs before and after CNO application. N = 3, n = 11. **(F)** Experimental timeline. **(G)** Coronal schematic showing DIO-hM4D(G_i_)-mCherry or DIO-mCherry injection into the dorsolateral striatum (DLS) with local cannula infusion of clozapine-N-oxide (CNO) or saline. FM4-64 dye infused prior to sacrifice. (H) Left, Coronal section showing loss of TH immunoreactivity in the ipsilesional hemisphere, expression of DIO-hM4D(G_i_)-mCherry (red), and infusion radius (FM4-64x, blue). The overlap is visible as a magenta color. Scale bar is 1 mm. *Right,* Schematic depicting infusion sites across animals. **(I)** Mean total abnormal involuntary movement (AIM) score following IP injection of CNO and IP injection of an individually titrated, sub-dyskinetic dose of levodopa (0.75 – 2 mg/kg) in animals expressing DIO-hM4D(G_i_)-mCherry or DIO-mCherry. Values represent the mean across animals at each time point. 2-Way RM ANOVA: p = 0.0059. **(J)** No dyskinesia was observed following IP injection of CNO and IP injection of saline. **(K)** Mean total AIM score following local striatal infusion of CNO and IP injection of the same individually titrated, sub-dyskinetic dose of levodopa (0.75 – 2 mg/kg). Values represent the mean across animals at each time point. 2-Way RM ANOVA: p < 0.0001. **(L)** No dyskinesia was observed following local striatal infusion of saline and IP injection of a low-dose of levodopa hM4D(G_i_): N = 14, mCherry: N = 12. N = animals. Data shown as mean ± SEM. A portion of the slice validation presented in (B) – (C) is used in another manuscript (*53*).

To validate the effects of CNO on basal ganglia activity *in vivo*, we performed freely moving electrophysiology in the main downstream target of D2-MSNs, the GPe (Figure S6A-C). We hypothesized that local striatal activation of hM4D(G_i_) through an infusion cannula in the DLS would selectively modulate local synaptic connections, without changing the intrinsic excitability of D2-MSNs, and thus that GPe firing rates would be unchanged. We found that neither local DLS infusion of saline or CNO evoked a significant change in GPe multi-unit activity (Figure S6D-I). As a positive control, in separate sessions we administered saline or CNO systemically, via IP injection (Figure S6J-O). As expected, we observed increased GPe activity with IP CNO (Figure S6K,M,O). These findings indicate that local striatal activation of G_i_-DREADD in D2-MSNs can alter MSN-MSN lateral inhibition, without altering the main synaptic output of D2-MSNs to the GPe.

Having validated this strategy, we tested the role of striatal lateral inhibition in mediating LID. Unilateral 6-OHDA-treated A2a-Cre mice were injected with either a Cre-dependent inhibitory DREADD (DIO-hM4D(G_i_)-mCherry) or an inert fluorophore control (DIO-mCherry) into the ipsilateral DLS (Figure 4F). After a period of recovery, animals began daily levodopa treatments. Later, an infusion cannula was implanted into the DLS to allow for local striatal delivery of the ligand CNO (or saline). At the end of all experiments, a lipophilic dye (FM4-64X) was infused just prior to sacrifice and used to confirm patency of the cannula in postmortem tissue (Figure 4G-H).

To assess the role of D2-mediated lateral inhibition in LID, we first sought to identify a low dose of levodopa (0.75-2 mg/kg) that increased movement (alleviated bradykinesia) but did not evoke dyskinesia. We anticipated we would be able to detect changes in the threshold for dyskinesia at this dose. Prior work suggests that chemogenetic stimulation of D2-MSNs abolishes the therapeutic response to levodopa in parkinsonian mice (*47*). To investigate how chemogenetic inhibition of D2-MSNs impacts dyskinesia expression, we delivered an IP injection of CNO followed by an IP injection of low-dose levodopa or saline. As predicted, a reduction of D2-MSN-mediated inhibition with CNO evoked dyskinesia (Figure 4I), while CNO paired with saline did not (Figure 4J).

While systemic administration of CNO in hM4D(G_i_)-expressing A2a-Cre animals provides further evidence for the role of D2-MSNs in shaping LID, this manipulation does not isolate the contribution of striatal lateral connections, as hM4D(G_i_)-positive terminals are present in both the striatum and the GPe. To determine if suppression of just the D2-MSN-mediated lateral inhibition lowers the threshold for LID, we infused CNO in the DLS, followed by an IP injection of a low-dose of levodopa. Under these conditions, hM4D(G_i_)-expressing mice became dyskinetic, while mCherry controls did not (Figure 4K). We performed several additional control experiments: animals expressing hM4D(G_i_) or mCherry did not develop abnormal involuntary movements to any other treatment pairing (striatal infusion/IP injection): saline/levodopa (Figure 4L), CNO/saline (Figure S7A), or saline/saline (Figure S7B). These *in vivo* experiments suggest that loss of D2-MSN-mediated lateral inhibition, combined with acute dopamine signaling, can contribute to LID.

## DISCUSSION

Despite a long-standing appreciation of striatal MSN-MSN connections and growing evidence for their functional significance, their contribution to motor dysfunction remains incompletely understood. Here, we used *ex vivo* and *in vivo* techniques to investigate how these connections change across healthy and disease states. Our overarching hypothesis was that these connections help coordinate the activity of functional striatal MSN ensembles involved in regulating motor output, and that disruption of these connections might contribute to involuntary movements.

In *ex vivo* brain slices, paired recordings across MSN-MSN subtypes provides critical information about the rate of connectivity and the strength of unitary connections (*19*). However, this approach is low throughput: depending on the connection type, up to 94% of MSN-MSN pairs lack a 1:1 connection (*19*). To overcome this obstacle, we employed an optogenetic approach (*30*) that allowed us to more comprehensively assess all MSN-MSN connection types. Using this technique, we reproduced the finding from paired recordings that MSN-MSN connections are asymmetric, with D2- to D1-MSN connections being the strongest overall. A limitation of the optical approach is that it yields a single number (compound inhibitory postsynaptic response amplitude), which likely reflects both the size of unitary responses and connection probability. Despite this limitation, our findings suggest the optical approach is an effective and efficient method for measuring the overall strength of MSN-MSN lateral inhibition.

We next sought to examine how MSN-MSN lateral inhibition is changed across disease states. A previous study using paired recordings found decreased strength of lateral inhibition onto D1-MSNs in the parkinsonian state, with no detectable D1-D1 connections (*19*). Consistent with these findings, we observed a decrease in the strength of D2-D1 oIPSC amplitudes. Importantly, the optical approach allowed us to record functional D1-D1 oIPSCs from parkinsonian animals. This finding suggests that although there is a marked reduction in the rate of D1-D1 connectivity, a modest network of D1-D1 connections remains in the parkinsonian state. Although our experiments focused on the DLS, similar adaptations in MSN-MSN lateral inhibition may occur in other striatal regions. Whether these changes differ across striatal subdivisions and contribute to distinct aspects of motor dysfunction remains to be determined.

High striatal firing rates have been observed in LID across several animal models (*6*, *12*, *33*, *35*, *36*). Further, movement-associated D1-MSN ensembles are larger in parkinsonian animals acutely treated with levodopa (*13*), and dyskinesia is associated with specific hyperactive D1- and D2-MSN populations (*11*). These changes in striatal activity likely reflect multiple cellular and circuit adaptations, including well-established hypersensitivity of D1-receptor signaling and alterations in excitatory synaptic transmission. However, we do not understand the microcircuit mechanisms which may lead to the recruitment of large and hyperactive movement-related ensembles in LID. We therefore tested whether changes in MSN-MSN lateral inhibition might represent an additional synaptic mechanism contributing to dyskinesia. Consistent with this hypothesis, we observed a restoration in D2-D1 inhibition following chronic levodopa treatment. This restoration may represent a compensatory response to repeated changes in striatal activity during dopamine replacement therapy, consistent with previously observed adaptations in the intrinsic properties of MSNs in the same mouse model (*34*). Recent work has demonstrated that chronic levodopa treatment induced GluN2B-dependent plasticity at glutamatergic synapses onto D2-MSNs during the OFF state, contributing to subsequent dyskinesia (*48*). Whether analogous adaptations occur at inhibitory synapses formed by D2-MSNs, potentially contributing to the restoration of D2-D1 lateral inhibition, remains unknown. Such adaptations could involve the number of functional synaptic connections, presynaptic neurotransmitter release, or postsynaptic responsiveness, although our measurements of compound oIPSCs cannot distinguish among these possibilities.

LID only occurs when levodopa is administered; our initial slice experiments did not capture this acute period of elevated dopamine signaling. To isolate the effects of acute D_2_R-activation on lateral inhibition in *ex vivo* brain slices, we applied the D_2_R-agonist quinpirole. Prior studies indicate that MSN-MSN lateral connections are acutely regulated by dopamine (*38*, *40*) and D2-D1 connections specifically are inhibited by the D_2_R agonist quinpirole (*30*, *38*). We replicated this finding in the DLS of healthy animals and discovered that D2-D1 connections are also inhibited by quinpirole in parkinsonian animals and parkinsonian animals chronically treated with levodopa. In the LID state, a dopamine-mediated reduction in D2-D1 lateral inhibition would be predicted to facilitate increases in both D1-MSN firing rates and ensemble sizes. This interpretation is consistent with recent observations that D1-MSNs become hyperexcitable during the LID ON-state (*14*) and that dyskinetic behaviors are associated with specific hyperactive D1-MSN populations (*11*). This proposed disinhibitory mechanism may act alongside other well-established changes like the hypersensitivity of D_1_R signaling following dopamine depletion (*49–51*), potentially contributing to D1-MSN overactivity during LID.

Synaptic transmission between MSNs has been described in organotypic cultures (*52*), the nucleus accumbens (*30*, *38*, *53*), and in the dorsal striatum (*19*, *52–54*). However, these studies do not directly establish the contribution of local MSN-MSN inhibitory synaptic transmission to motor dysfunction *in vivo*. A prior study in the nucleus accumbens found that both D2-D1 connections and cocaine-induced locomotor hyperactivity were dependent on dopamine D_2_R, but could not directly show that changes in lateral inhibition caused hyperactivity (*30*). One obstacle to identifying the behavioral role of MSN-MSN connections is that MSNs may regulate circuit function and behavior via local GABA release in the striatum and/or via their connections to downstream basal ganglia nuclei. Chemogenetic or optogenetic activation of D2-MSNs reduces the behavioral response to levodopa, causally linking D2-MSNs to LID (*47*, *55*). From this literature, it is unclear whether D2-MSNs regulate LID via local striatal connections, their output to GPe, or both. We took advantage of the lack of GIRK effectors (*45*, *46*) to inhibit MSN synaptic outputs in the striatum without affecting their overall firing or synaptic output downstream. This approach allowed us to investigate the contribution of local D2-MSN-mediated inhibition to dyskinesia expression *in vivo*. We found that disruption of D2-mediated lateral connections lowers the threshold for the expression of dyskinesia and explains part of the overall effect of inhibiting D2-MSNs in dyskinesia. A caveat to our chemogenetic approach is that the manipulation targets a pre-synaptic cell type. Future work could utilize alternative techniques, such as drugs acutely restricted by tethering (DART)(*56*) to achieve post-synaptic specificity.

Interestingly, systemic and local striatal CNO administration produced similar peak AIM scores, although the dyskinetic response was more prolonged following systemic administration (Figure 4). However, these administration methods differ in their kinetics and spatial extent, and the effective concentration of CNO achieved in the striatum cannot be directly compared. Although systemic CNO administration increased GPe activity whereas local infusion did not (Figure S6), these differences preclude determining whether the more prolonged dyskinetic response following systemic administration reflects the recruitment of downstream D2-MSN projections or differences in CNO exposure. Together, these findings highlight the importance of D2-MSN-mediated inhibition in regulating dyskinesia and the potential contributions of both local and downstream circuitry.

More broadly, although D1-MSNs have been a focus of recent studies in LID, a growing body of evidence points to an important role for D2-MSNs. In retrospect, part of this functional impact might be explained by D2-MSN-mediated lateral inhibition. Emerging evidence suggests that D2-MSNs undergo plastic changes with chronic dopamine replacement and selective disruption of D2 receptor signaling strongly alters dyskinetic output (*34*, *48*, *57*, *58*). Studies of D_2_R agonists underscore the importance of D2-mediated inhibition in regulating dyskinesia-related activity. For example, quinpirole and pramipexole enhance D1-MSN activation in rodent models of PD/LID (*12*, *47*, *55*, *59–61*). Together with the current findings, these studies point to a multifaceted role of D2-MSN activation in levodopa treatment, involving both (1) disinhibition of LID-associated ensembles in the GPe (*62*), and (2) disinhibition of D1-MSN activity locally in the striatum.

In addition to lateral inhibition from neighboring MSNs, MSNs also receive inhibitory input from several interneuron populations, including parvalbumin-expressing fast-spiking interneurons and somatostatin/neuropeptide Y-expressing interneurons (*16*). Previous studies have demonstrated that fast-spiking interneuron connectivity is altered following dopamine depletion and that disruption of fast-spiking interneuron activity can contribute to dyskinetic movements (*17*). Although our experiments specifically examined D2-MSN-mediated lateral inhibition, these findings should be considered alongside broader changes in striatal inhibitory microcircuits that may contribute to LID.

The proposed role of striatal lateral inhibition in shaping MSN ensemble activity parallels well-established functions of lateral inhibition in other brain regions. In sensory systems, lateral inhibition plays a role in improving sensory perception by shaping the pattern of neural activity evoked by sensory stimuli (*22–29*). Lateral inhibition sharpens stimulus representations by suppressing surrounding activity and enhancing contrast. Analogous to this role, lateral inhibition in the striatum may serve to refine or constrain competing motor representations. Although our experiments were conducted in disease models, the observed modulation of motor output by lateral inhibition suggests that this circuit motif may have broader relevance for regulating MSN ensemble activity in the intact brain (*10*, *63*, *64*). Whether lateral inhibition directly shapes MSN ensemble organization or selection among competing actions during normal motor behavior remains to be determined. The extensive local connectivity of MSNs provides an anatomical and physiological basis for this proposed role. MSNs have elaborate axonal arborizations within the striatum, extending up to 0.5 mm (*2*, *65*). Although MSN-mediated connections have small unitary connection strengths, they form at rates up to 36%, and comprise >90% of all striatal neurons (*1*, *19*). The sheer size of this inhibitory plexus and its spatial organization may have strong implications for shaping striatal output.

## MATERIALS AND METHODS

## RESOURCE AVAILABILITY

### Lead contact

Requests for additional information, resources, or reagents should be directed to and will be fulfilled by the Lead contact, Alexandra Nelson.

### Materials availability

This study did not generate any new unique materials.

## EXPERIMENTAL MODEL AND STUDY PARTICIPANT DETAILS

### Animals

All animal procedures were approved by the University of California San Francisco Institutional Animal Care and Use Committee (IACUC; AN202979-00C). Animals were housed under a 12h light/dark cycle with access to food and water ad libitum. For all experiments, both male and female C57Bl/6J mice, aged 3-12 months, were used. For *ex vivo* electrophysiology experiments, A2a-Cre;D1-tdTomato, D1-Cre;D1-tdTomato, and A2a-Cre;D2-GFP mice positive for both transgenes were used. For *in vivo* cannula infusion experiments, A2a-Cre animals were used. Age-matched littermates were randomly assigned to experimental and control groups.

## METHOD DETAILS

### Surgical Procedures

A detailed surgical protocol can be found at (dx.doi.org/10.17504/protocols.io.b9kxr4xn). Briefly, three- to six-month old mice were anesthetized with a combination of ketamine/xylazine (40/10 mg/kg IP) and inhaled isoflurane (1%). After placement in the stereotaxic frame (Kopf Instruments), the scalp was opened, and a mounted drill was used to create a hole over the left medial forebrain bundle (MFB). Using a 33-gauge needle (WPI) and Micro4 pump (WPI), 1 μL of 6-hydroxydopamine (6-OHDA, Sigma-Aldrich, 5 μg/μl in normal saline) was injected unilaterally into the MFB (-1.0 AP, -1.0 ML, -4.9 DV) at a rate of 0.2 μl/min. To reduce toxicity to other monoaminergic axons, desipramine (Sigma-Aldrich, 25mg/kg IP) was administered 30 min prior to 6-OHDA injection. Control animals were injected with 1 μL of normal saline in the MFB using the same parameters. After the operation, animals received daily saline injections and were provided high-fat dietary supplements (Diet-Gel, peanut butter, forage mix) for a 1-2 week recovery period.

Viral injections were performed during the same surgical procedure as the MFB injection described above. A mounted drill was used to create a hole over the left dorsolateral striatum (DLS). Using a 33-gauge needle (WPI) and a Micro4 pump (WPI), 1 μL of double-floxed inverse open reading frame (DIO) constructs were used to express virus in a pathway-specific manner. For *ex vivo* electrophysiology experiments, mice were injected with 1 μL AAV5-EF1a-DIO-hChR2-(H134R)-eYFP-wpre-HGH (1:3 dilution, Penn) into the left DLS (+0.8 AP, -2.3 ML, -2.5 DV). For chemogenetic inhibition experiments, mice were injected with 1 μL of AAV5-hSyn-DIO-hM4D(G_i_)-mCherry (1:2 dilution, Addgene), or AAV5-hSyn-DIO-mCherry (undiluted, UNC) into the left DLS. For chemogenetic validation experiments measuring D2-D1 synaptic transmission, A2a-Cre x D2-GFP mice were injected with a mixture of AAVs encoding ChR2-eYFP and hM4D(G_i_)-mCherry (1 μL total injection volume). For separate current-clamp experiments measuring D2-MSN intrinsic excitability, A2a-Cre mice were injected with AAV encoding hM4D(G_i_)-mCherry.

For animals receiving local striatal infusions, the scalp was reopened 3–4 weeks after the initial surgeries to implant guide cannula assemblies in the DLS (5 mm pedestal with 0.05 mm protrusion; Protech International). Cannulas were secured in place with dental cement (Metabond) and dental acrylic (Ortho-Jet or Henry Schein).

For *in vivo* electrophysiology experiments, the scalp was reopened 3–4 weeks after the initial surgeries to implant guide cannula assemblies in the DLS (5 mm pedestal with 0.05 mm protrusion; Protech International) and 32-channel arrays (Innovative Neurophysiology) in the GPe. Three additional holes were drilled for two skull screws (FST) and a ground wire. The array was slowly lowered through the craniectomy into the GPe (AP: -0.4, ML: -1.9, DV: -4.0). The cannula assemblies were implanted at a 20° off the vertical, such that the cannula base pointed anteriorly (coordinates of the burr hole for cannula insertion AP: +1.7, ML: -2.4). The guide cannula tip was advanced along the axis to a final DV of -2.5 mm.

### Behavior

Details of the behavioral assessments can be found at (dx.doi.org/10.17504/protocols.io.b9ksr4we). Gross movement was quantified across several metrics, using video-tracking software (Noldus Ethovision) of the animals in an open-field arena (25 cm diameter cylinder). Velocity, distance traveled, and rotational behavior were measured. Rotation rate was calculated in 60 s bins by subtracting total ipsilesional rotations from total contralesional rotations.

After a 3-week post-surgical baseline period, animals began daily treatment. Parkinsonian animals receiving levodopa developed robust levodopa-induced dyskinesia (LID). Dyskinesia was quantified using the Abnormal Involuntary Movement score (AIMs)(*66*), a standardized metric that takes into account dyskinesia across axial, limb, and orolingual body segments. As described by Cenci and Lunblad (2007), dyskinesia severity in each body segment was quantified during 60 s observation windows as follows: 0 = normal movement, 1 = abnormal movement for < 50% of the time, 2 = abnormal movement for > 50% of the time, 3 = abnormal movement for 100% of the time, but can be interrupted, and 4 = continuous, uninterruptable abnormal movement. The total AIM score represents the summation of the AIM score for each body segment (axial, limb, orolingual), for a maximum possible score of 12. For weekly scoring sessions, blinded experimenters rated AIMs every 20 m for a 2 h period. For cannula infusion experiments, dyskinesia was monitored every other minute.

### Pharmacology

Details for the preparation of pharmacological agents can be found at (dx.doi.org/10.17504/protocols.io.e6nvw1nbzlmk/v2). For *in vivo* experiments, levodopa (Sigma-Aldrich) was co-administered with benserazide (Sigma-Aldrich) and prepared in normal saline. For *ex vivo* electrophysiology experiments, 5–10 mg/kg levodopa and 2.5–10 mg/kg benserazide were used. Levodopa was given via IP injection 5–7 days per week. Daily levodopa injections continued for 3 or more weeks prior to experiments. Clozapine N-Oxide (CNO) was dissolved in normal saline. CNO was delivered via local striatal infusion (10 μM) or via IP injection (1 mg/kg).

Prior to intracranial infusion experiments, a sub-dyskinetic dose of levodopa (0.75-2 mg/kg) was determined for each mouse. For the dose to be deemed sub-dyskinetic, the animal needed to display a prokinetic effect (contra-lesional rotational bias, increased velocity and distance traveled) with no dyskinesia (AIM score = 0). After the sub-dyskinetic doses were determined, the infusion experiments commenced. For infusion experiments, the experimenter was blinded to the viral construct injected in the mice. After a baseline period in the open-field arena, the dummy cannula was removed, and the infusion cannula was screwed into the guide. On interleaved days, CNO (10 μM) and saline were infused using a Hamilton syringe pump (WPI) at a rate of 0.2 μl/min for a total volume of 2 μl. The experimenter was also blinded to the infusion liquid. The infusion cannula was removed 10 min following the conclusion of the infusion and the dummy cannula was restored. Following a 10 min infusion baseline period, the mouse received an IP injection (sub-dyskinetic dose of levodopa or normal saline, delivered on interleaved days). Following IP injection, AIMs were monitored for 60 s every-other minute for 60 min. Infusion experiments were performed a minimum of 24 h apart.

To verify the placement and patency of the infusion cannulas, as well as the spread of the drug infusion, a lipophilic fluorescent dye (FM4-64X, Thermo-Fisher Scientific) was infused before sacrifice. The dye was infused using the same experimental parameters used for CNO infusion (2 μl at 0.2 μl/min). Fifteen min after the conclusion of the infusion, the animal was terminally anesthetized and transcardially perfused for immunohistochemistry.

For *ex vivo* experiments, picrotoxin (Sigma-Aldrich) was dissolved in warm water at 5 mM and added to ACSF for a final concentration of 50 μM. Quinpirole (Tocris) was dissolved in water and added to ACSF for a final concentration of 1 μM. CNO (Tocris) was dissolved in water at 10 mM and added to ACSF for final concentrations of 1 μM or 10 μM.

### *Ex vivo* electrophysiology

A detailed protocol for *ex vivo* electrophysiology can be found at (dx.doi.org/10.17504/protocols.io.b9uir6ue). To prepare acute brain slices, mice were anesthetized with ketamine/xylazine and transcardially perfused with 95%O_2_/5%CO_2_ oxygenated, ice-cold glycerol-based artificial cerebrospinal fluid (ACSF) containing (in mM): 250 glycerol, 2.5 KCl, 1.2 NaH_2_PO_4_, 10 HEPES, 21 NaHCO_3_, 5 D-Glucose, 2 MgCl_2_, 2 CaCl_2_. Following decapitation, brains were dissected and sequential coronal slices (275 μm) containing the striatum were collected with a vibratome (Leica). Slices were immediately transferred to a holding chamber containing warm (33°–34°C), carbogenated ACSF containing (in mM): 125 NaCl, 26 NaHCO_3_, 2.5 KCl, 1.25 NaH_2_PO_4_, 12.5 D-Glucose, 1 MgCl_2_, 2 CaCl_2_. Slices were incubated for 30 min and then held at room temperature (22°–24°C) until used for recording.

For all recordings, slices were transferred to a recording chamber superfused (∼2 mL/min) with carbogenated ACSF (31°–33°C). Medium spiny neurons were targeted for recordings using differential interference contrast (DIC) optics on an Olympus BX 51 WIF microscope. In Drd1a-tdTomato mice (Figures 1-3), direct pathway neurons were identified by their tdTomato-positive somata. Conversely, indirect pathway neurons were identified by their medium-sized tdTomato-negative somata. In D2-GFP mice (Figure 4A-E), indirect pathway neurons were identified by their GFP-positive somata and direct pathway neurons were identified by their GFP-negative, medium-sized somata. Fluorescent-negative neurons with interneuron-like physiological properties were excluded from the dataset. In A2a-Cre x D2-GFP mice used for voltage-clamp validation experiments (Figure 4B-C), putative D1-MSNs were identified by their medium-sized somata and absence of both green fluorescence (D2-GFP/ChR2-eYFP) and red fluorescence (hm4D(G_i_)-mCherry). In separate current-clamp experiments (Figure 4D-E), D2-MSNs were identified by hM4D(G_i_)-mCherry expression in A2a-Cre mice. Fluorescence-negative neurons with interneuron-like physiological properties were excluded from the dataset.

Medium spiny neurons were patched in the whole-cell configuration using borosilicate glass electrodes (2–5 MΩ). For voltage-clamp experiments, electrodes were filled with a cesium methanesulfonate-based internal solution with high chloride containing (in mM): 120 CsCl, 15 CsMESO_3_, 8 NaCl, 0.5 EGTA, 10 HEPES, 5 QX-314, pH 7.3. For current-clamp experiments, electrodes were filled using a K-based internal solution containing (in mM): 130 KMeSO_3_, 10 NaCl, 2 MgCl_2_, 0.16 CaCl_2_, 0.5 EGTA, 10 HEPES, 2 MgATP, 0.3 NaGTP, pH 7.3). After whole-cell break in, cells were given at least 5 min to dialyze internal solution before recording. Whole-cell recordings were made using a MultiClamp 700B amplifier (Molecular Devices) and digitized with an ITC-18 A/D board (HEKA). Data were acquired using Igor Pro 6.0 software (Wavemetrics) and custom acquisition routines (mafPC, courtesy of M.A. Xu-Friedman).

Inhibitory currents were optically evoked using 3 ms pulses of 473 nm light, ranging in power from 0.5-4 mW delivered by a TTL-controlled LED (Olympus) passed through a GFP filter (Chroma). Cells were voltage-clamped at -70 mV. Series resistance was monitored throughout the experiment, and cells were removed from analysis if the change in access was >20% from the baseline period. To determine if D1-MSNs had an intrinsic ChR2 current (in Drd1a-Cre animals injected with DIO-ChR2-eYFP), a 500 ms test pulse was delivered. Neurons with resulting 500 ms inward currents and/or response latency of <2 ms were deemed to have an intrinsic ChR2 current and excluded from further analysis. The same exclusion criteria were applied to D2-MSNs in Adora2a-Cre animals injected with DIO-ChR2-eYFP.

### *In vivo* electrophysiology

A detailed protocol for *in vivo* electrophysiology can be found at (dx.doi.org/10.17504/protocols.io.36wgq641ylk5/v1). Briefly, mice were habituated to the open field, tethering, IP injections, and cannula infusions. The animal’s gross behavior was recorded (IC Capture, ImagingSource) and analyzed using video tracking software (Noldus Ethovsion). Recordings consisted of a 30 m baseline period followed by IP injection or cannula infusion. Electrophysiological recordings continued for at least 60 m post-manipulation. Behavioral measurements were synchronized with simultaneous electrophysiological recordings via TTL pulses and recorded by the electrophysiology system.

Recordings were performed with a 32-channel fixed electrode array in the globus pallidus *pars externa* (AP: -0.4, ML: -1.9, DV: -4.0). Signals were recorded at 30 kHz using an Intan system (RHD2000 Interface Board, Intan Technologies). Animals were placed in the open field and tethered by a multiplexed headstage cable attached to a low-torque electrical commutator (Doric). Cannula assemblies were implanted at a 20° angle in the DLS (coordinates zeroed from tip of dummy, AP: +1.7, ML: -2.4, DV: -2.5).

Single-unit (SU) and multi-unit (MU) activity was classified using automated spike sorting (KiloSort, https://github.com/cortex-lab/Kilosort) and manually curated using Phy (https://github.com/cortex-lab/phy). Single-unit activity was isolated for example traces and waveforms only. For MUA analysis, any single-unit and multi-unit activity detected on the same electrode were collapsed. Firing rate was calculated by averaging 1 m bins over the course of the experimental session. Modulation of the firing rate was determined by comparing firing rates pre- and post-manipulation, averaging over 30 m periods. Average multi-unit activity was calculated using firing rates normalized to baseline.

To verify the position of the electrode in GPe, mice were deeply anesthetized after the last recording session. Prior to transcardial infusion, electrolytic lesions were made to mark electrode tips using a solid-state, direct current Lesion Maker (Ugo Basile) by applying 100 μA for 5 s per microwire.

### Histology and microscopy

A detailed protocol for preparation of histological sections can be found at (dx.doi.org/10.17504/protocols.io.b9ubr6sn). After behavioral or chemogenetic experiments, mice were anesthetized with ketamine/xylazine and transcardially perfused with 4% paraformaldehyde (PFA) in phosphate buffer solution (PBS). Following perfusion, brains were dissected and post-fixed overnight in 4% PFA, then transferred to 30% sucrose and stored at 4°C for cryoprotection. Brains were then sliced coronally into 30 μm sections on a freezing microtome (Leica).

After *ex vivo* electrophysiology experiments, 275 μm slices were stored overnight in 4% PFA. The following day, slices were transferred to 30% sucrose and sub-sectioned into 50 μm sections on a freezing microtome (Leica).

For immunohistochemistry, the tissue was blocked in 3% normal donkey serum (NDS) and permeabilized with 0.1% Triton X-100 on a room-temperature (RT) shaker. Primary anti-bodies (Rabbit anti-TH, Pel-Freez Biologicals, 1:1000) were added to the NDS and tissue was incubated overnight on a 4°C shaker. Tissue was then incubated in secondary antibodies (Donkey anti-rabbit Alexa Fluor 488 or 594, 1:500) overnight (16 h) on a 4°C shaker before being washed and mounted (Vectashield Mounting Medium) onto glass slides for imaging. Images (4x, 10x, or 40x) were obtained using a Nikon 6D conventional widefield microscope or an Olympus Fluoview FV3000 confocal microscope. In 6-OHDA treated mice, the extent of dopamine depletion was confirmed by TH immunohistochemistry. Adequate expression of DIO constructs (ChR2-eYFP, eYFP, hM4D(G_i_)-mCherry, mCherry) was confirmed.

## QUANTIFICATION AND STATISTICAL ANALYSIS

Details can be found in Table S1. *Ex vivo* electrophysiology traces were processed in Igor Pro 6.3 (Wavemetrics). Statistical tests were performed using GraphPad Prism 10. All data are presented as the mean ± SEM, with “n” referring to the number of cells and “N” referring to the number of animals. Average amplitudes of oIPSCs collected using 1 mW light power were compared across MSN-MSN connection types (Figure 1G), as well as across treatment conditions (Figures 2G, 2I) using a nonparametric Kruskal-Wallis (KW) test. A post-hoc Dunn’s test for multiple comparisons was used. MSN-MSN oIPSC amplitudes following application of picrotoxin were compared using a paired, non-parametric Wilcoxon signed-rank (WSR) test (Figure S1). D2-D1 oIPSC amplitudes following application of quinpirole were also compared using a paired, non-parametric WSR for Ctrl, Park, and LD conditions (Figure 3A-C). The normalized oIPSC amplitude after application of quinpirole was compared across treatment conditions using a KW test (Figure 3A-C). A Friedman’s test with Dunn’s post-hoc tests were used to compare D2-D1 oIPSC amplitudes following application of quinpirole and sulpiride (Figure S4). A two-way repeated measures analysis of variance (2-way RM ANOVA) was used to compare AIMs score over time in systemic (IP injection) and infusion (cannula) chemogenetic inhibition experiments (Figures 4I, 4K). In *ex vivo* brain slices, a WSR test was used to compare measures of D2-MSN excitability following application of the ligand CNO. *In vivo*, a WSR test was used to compare multi-unit activity in the GPe following striatal infusion of saline (Figure S6H) or CNO (Figure S6I), as well as systemic injection of saline (Figure S6N) and CNO (Figure S6O).

## ACKNOWLEDGEMENTS

We thank Dr. Rodrigo Paz for invaluable assistance. We thank Drs. Massimo Scanziani, Kevin Bender, and Felice Dunn and all members of the Nelson Lab for helpful feedback on the project and manuscript. An earlier version of this manuscript was posted to BioRxiv on 12 October 2024 at doi: https://doi.org/10.1101/2024.10.11.617939.

## AUTHOR CONTRIBUTIONS

Conceptualization, E.L.T. and A.B.N.; Methodology, E.L.T., C.J.B.M., & A.B.N.; Investigation, E.L.T., O.K.B., L.S., C.J.B.M., A.E.G., S.S., and A.B.N.; Formal Analysis, E.L.T. and A.B.N.; Writing – Original Draft, E.L.T. and A.B.N.; Writing – Review & Editing, all authors; Visualization: E.L.T. and A.B.N.; Funding Acquisition, E.L.T. and A.B.N.; Supervision, A.B.N.

## DATA, CODE, AND MATERIALS AVAILABILITY

The data, code, protocols, and key lab materials used and generated in this study are listed in a Key Resource Table alongside their persistent identifiers at Zenodo (DOI: 10.5281/zenodo.24432115). Any additional information required to reanalyze the data reported in this paper is available from the lead contact upon request.

