## Supplementary Figures and Tables for "Striatal lateral inhibition regulates abnormal movements in a mouse model of levodopa-induced dyskinesia"

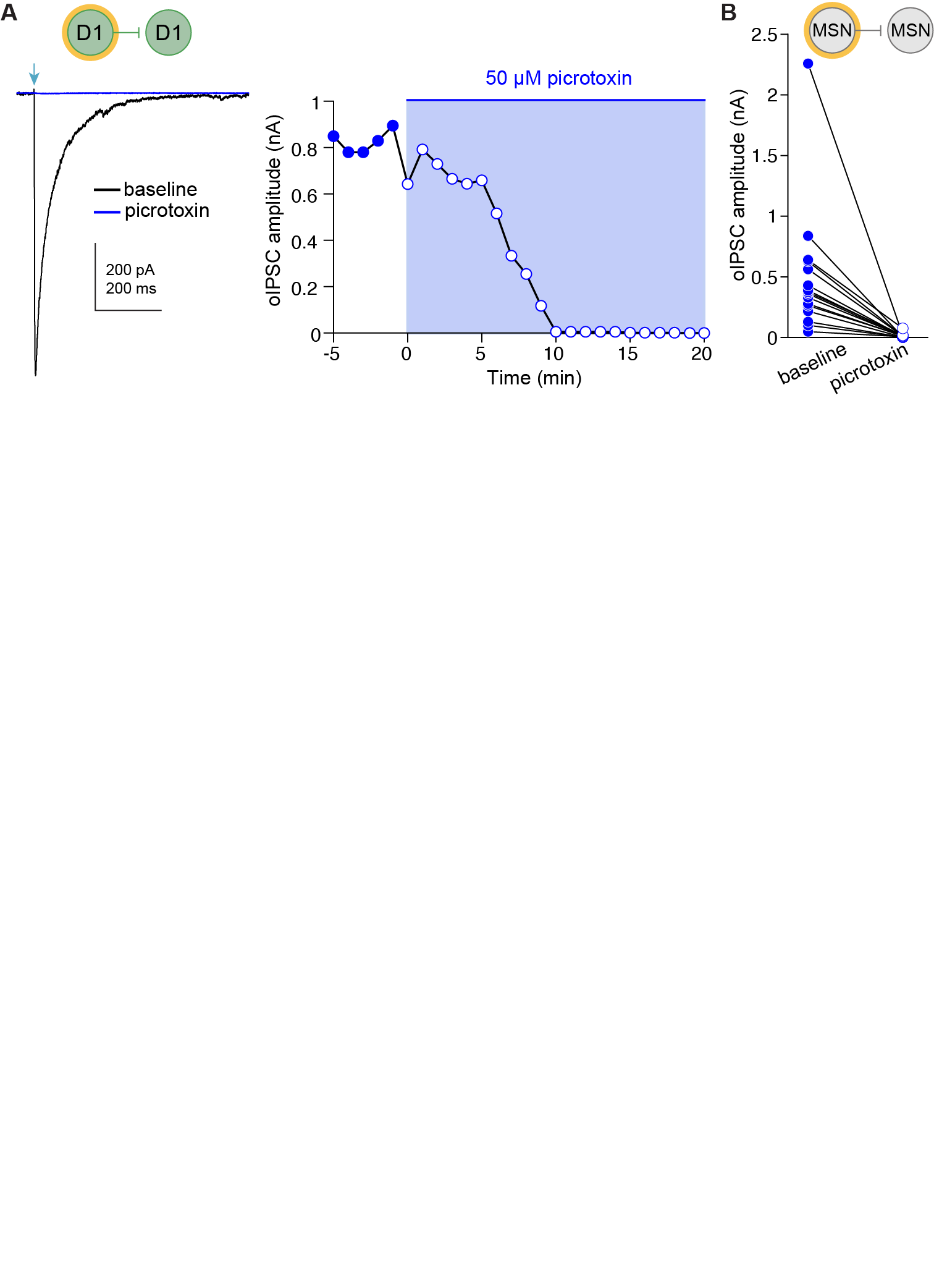

**Figure S1.** *MSN-MSN lateral connections are mediated by GABA_A_ receptors, related to Figure 1.*

(A) (*Left*) Representative example of D1-D1 oIPSC before and after application of 50 μM picrotoxin. (*Right*) Time course of the effect of picrotoxin on oIPSC amplitude.
(B) Summary of change in oIPSC amplitude in response to picrotoxin. MSN-MSN connection types are pooled. N = 13, n = 16. WSR: p < 0.0001. N = animals; n = cells. Data shown as mean ± SEM.

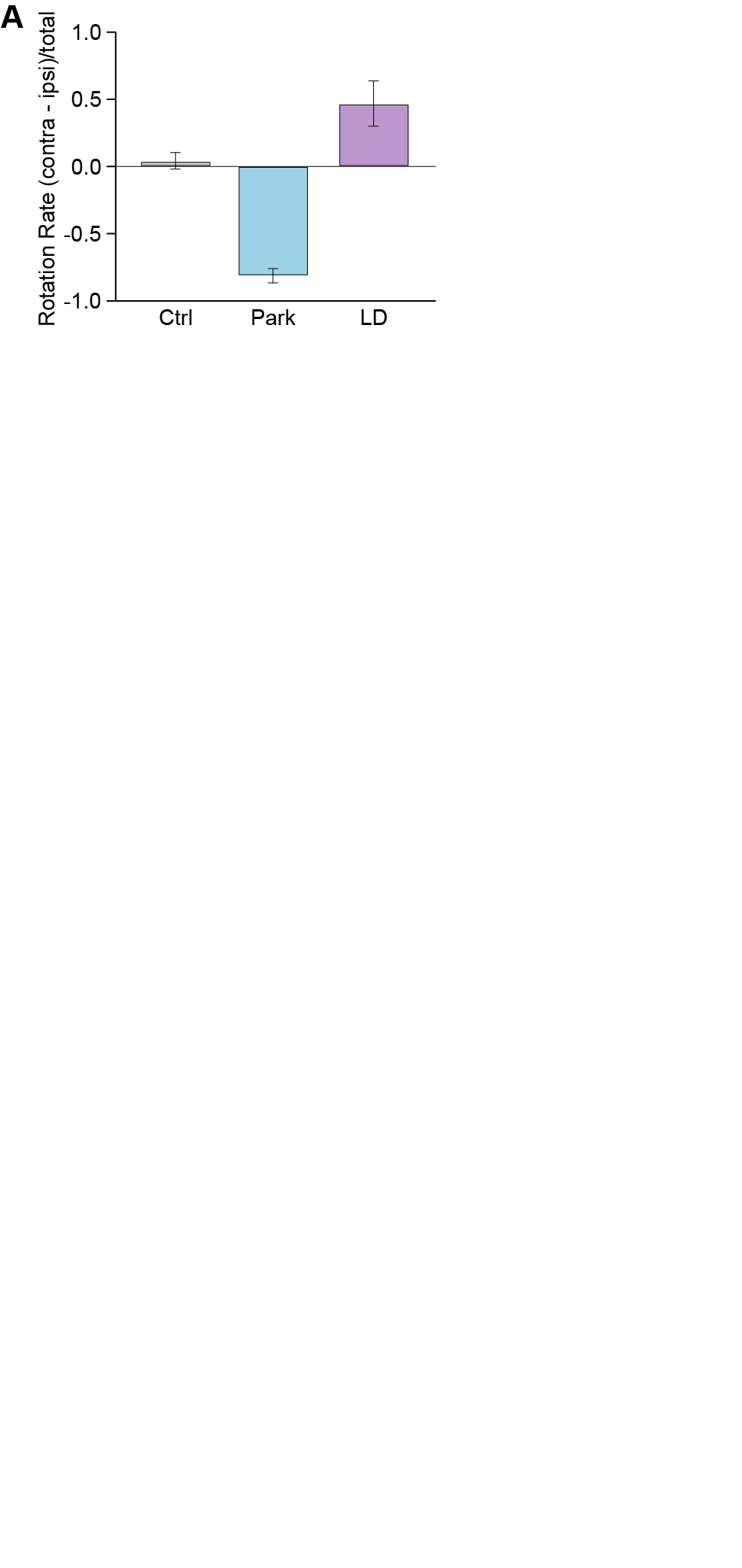

**Figure S2.** *Unilateral depletion of dopamine causes rotational bias, related to Figure 2.*

(A) Rotation rate in the open field over 10 minutes control (Ctrl), parkinsonian (Park and LD OFF medication), and parkinsonian mice treated with levodopa (LD ON medication). Rotation rate is calculated as number of ipsilesional rotations subtracted from the number of contralesional rotations divided by the total number of rotations. Ctrl: N = 9, Park: N = 14, LD: N = 8. N = animals. Data shown as mean ± SEM.

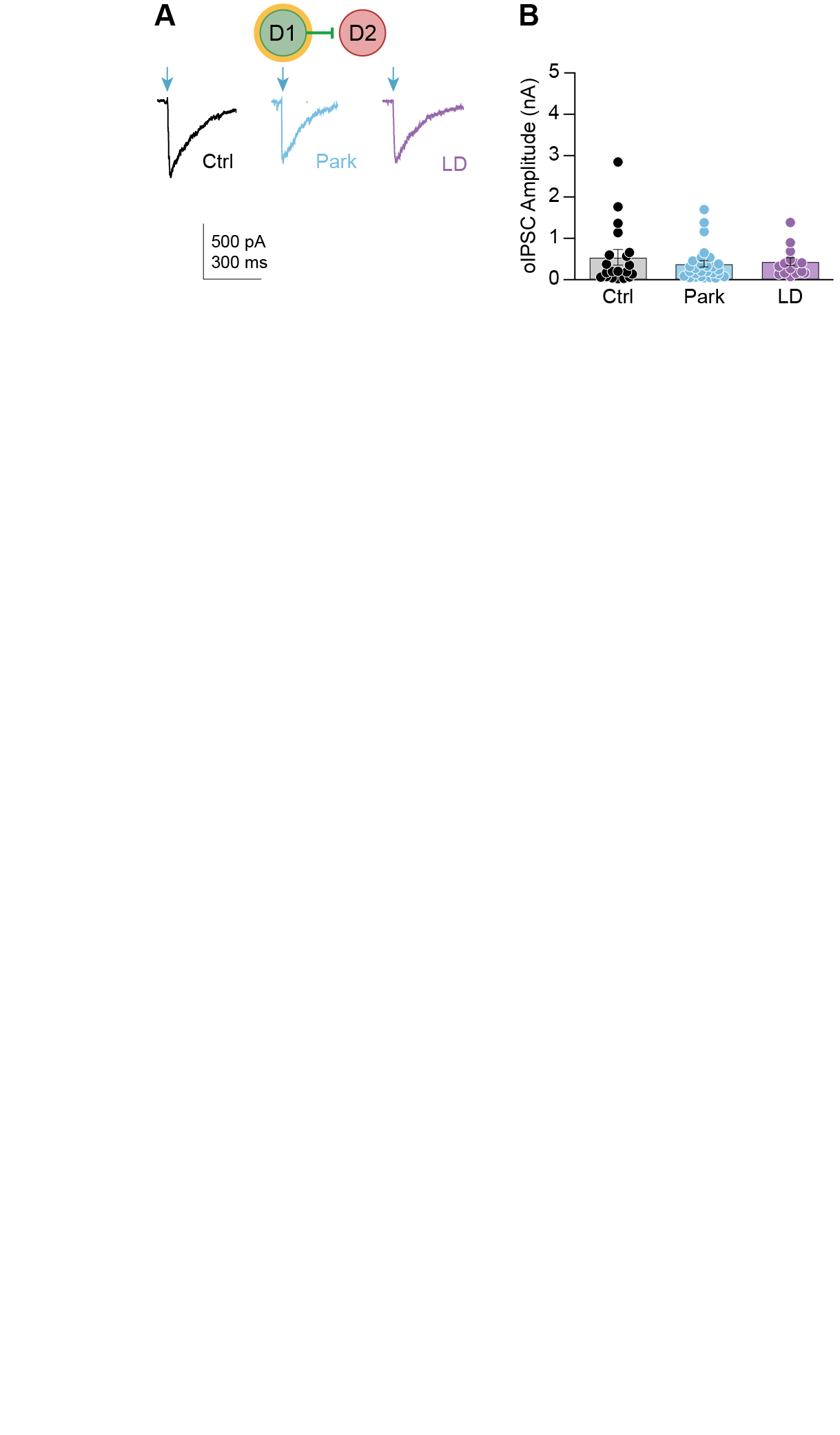

**Figure S3.** *D1-D2 synaptic responses do not differ across experimental groups, related to Figure 2.*

(A) Representative D1-D2 oIPSCs in animals belonging to Ctrl (*left*), Park (*middle*), and LD (*right*) groups.
(B) Average D1-D2 oIPSC amplitude in response to 1 mW optical stimulation for Ctrl (N = 9, n = 20), Park (N = 4, n = 30), and LD (N = 9, n = 16) groups. KW: Χ^2^ = 0.8926, p = 0.64. Each overlaid dot represents one cell. N = animals; n = cells. Data shown as mean ± SEM.

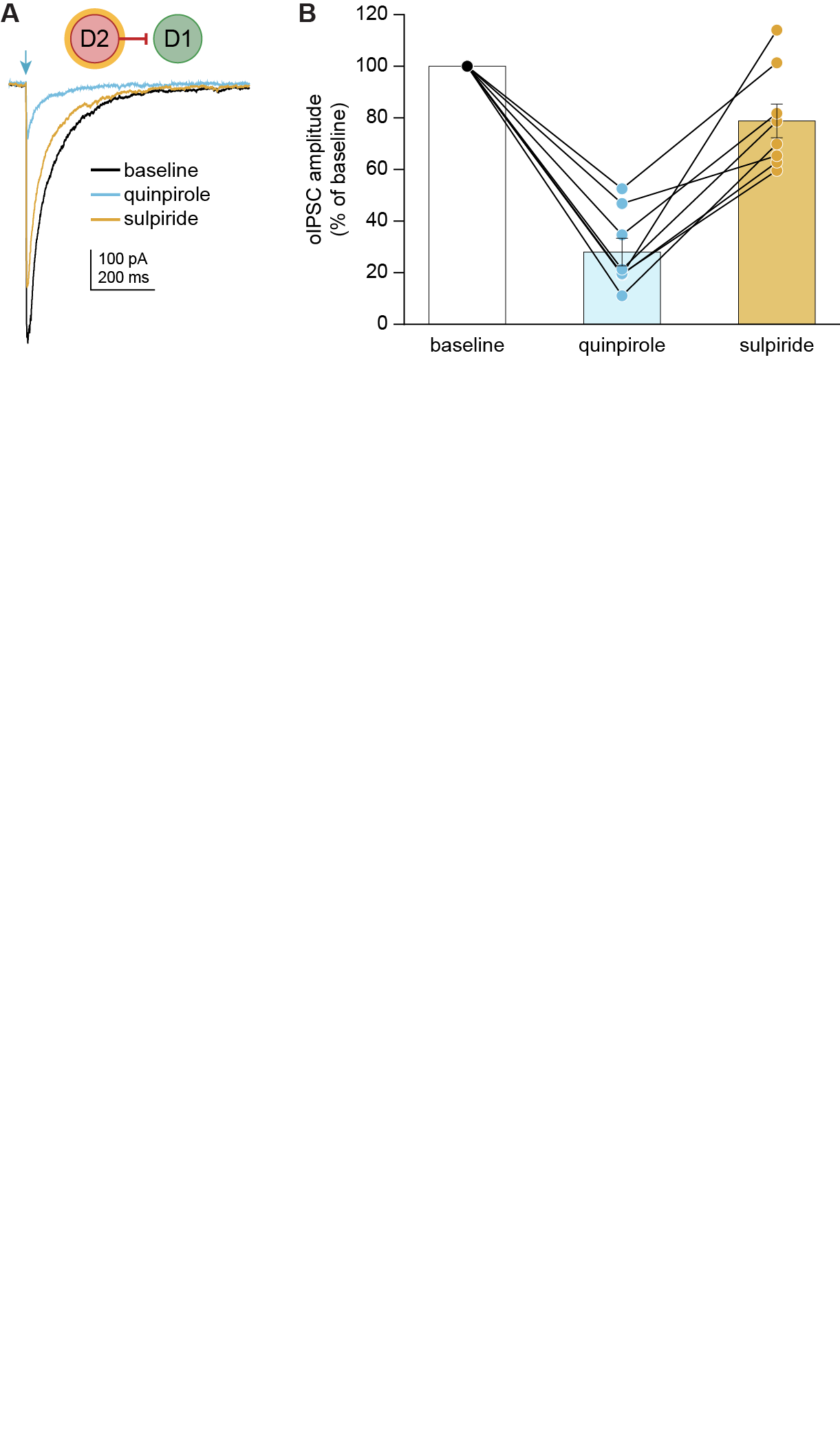

**Figure S4.** *D2-D1 oIPSCs are bidirectionally modulated by D_2_R agonist and antagonist, related to Figure 3.*

(A) Representative example of D2-D1 oIPSC and the effect of 1 μM quinpirole followed by 1 μM sulpiride in parkinsonian DLS.
(B) Summary of change in oIPSC amplitude in response to quinpirole followed by sulpiride. N = 3, n = 8. Friedman: Χ^2^ = 13, p = 0.0003. N = animals, n = cells. Data shown as mean ± SEM.

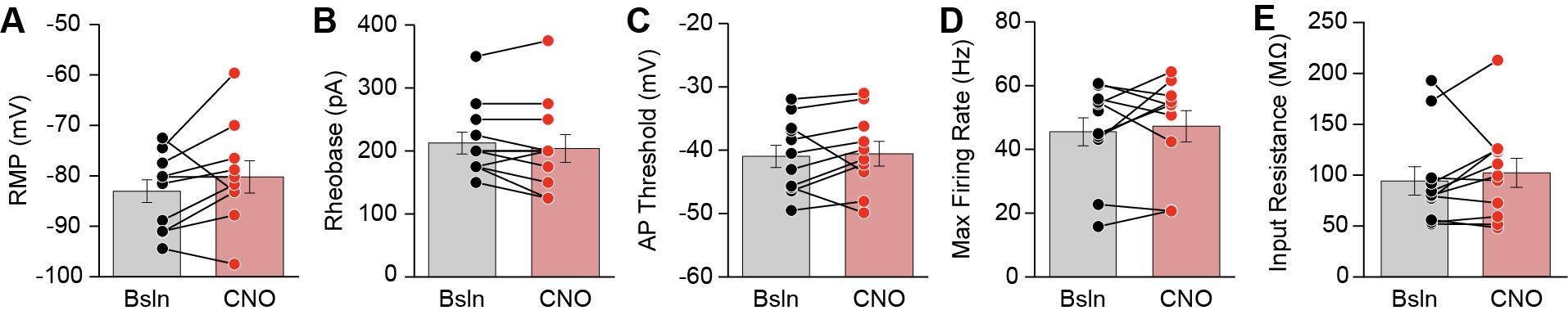

**Figure S5.** *CNO activation of hM4D(Gi) in D2-MSNs does not alter intrinsic membrane properties or excitability, related to Figure 4.*

(A) – **(E)** Ex vivo whole-cell current-clamp recordings from D2-MSNs in A2a-Cre;D2-GFP animals injected with AAV encoding hM4D(Gi)-mCherry. Membrane properties were compared before (Bsln) and after (CNO) application of 1 μM CNO. No change was observed for (A) resting membrane potential (mV, WSR: p = 0.13), (B) rheobase (pA, WSR: p = 0.36), (C) action potential threshold (mV, WSR: p = 0.32), (e) maximal firing rate (Hz, WSR: p = 0.56), or (E) input resistance (MΩ, WSR: p = 0.41) in D2-MSNs. N = 3, n = 11. Each dot represents one neuron. N = animals, n = cells. Data shown as mean ± SEM.

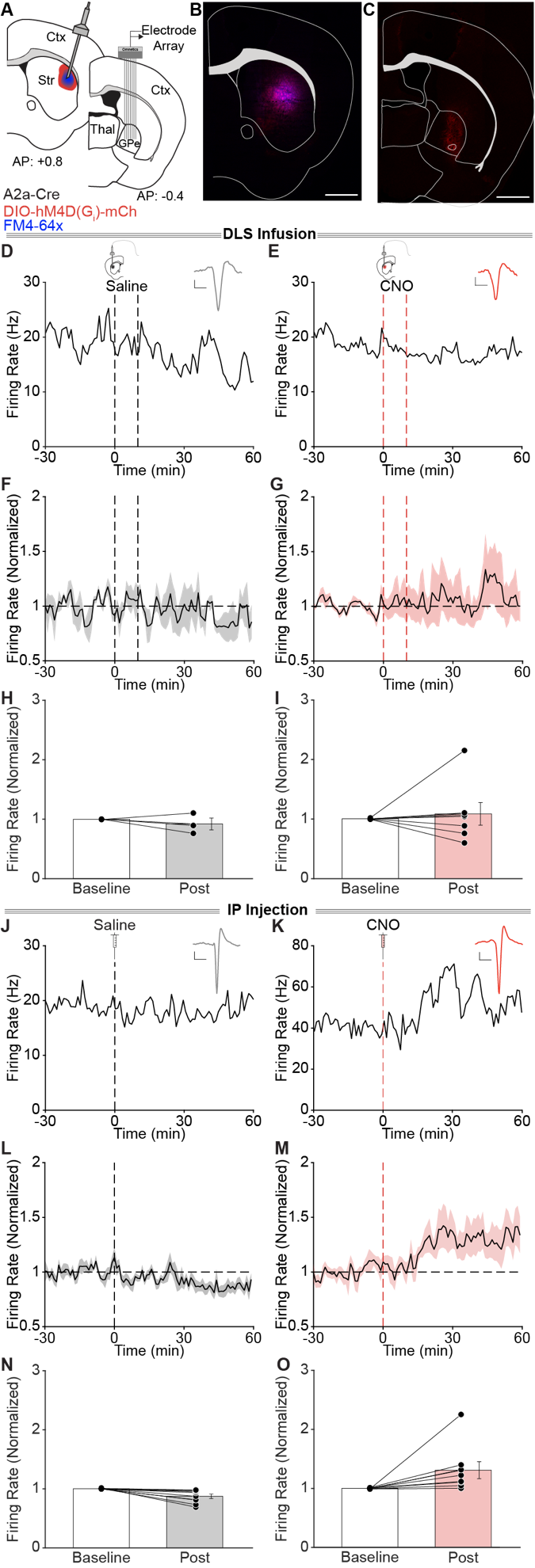

**Figure S6.** *Striatal CNO activation of hM4D(Gi) in D2-MSNs does not increase GPe activity in vivo, related to Figure 4.*

(A) – **(O)** *In vivo* validation of hM4D (Gi) in D2-MSNs. Parkinsonian, chronically levodopa-treated A2a-Cre mice were injected with AAV encoding Cre-dependent hM4D(Gi)-mCherry in the dorsolateral striatum (DLS), implanted with an infusion cannula in the DLS and multielectrode array in the globus pallidus *pars externa* (GPe).
(A) Coronal schematics depicting viral expression, infusion radius, and cannula placement in the DLS and 32-channel array placement in the GPe.
(B) Coronal section showing striatal expression of DIO-hM4D(G_i_)-mCherry (red), and infusion radius (FM4-64X, blue). The overlap is visible as magenta. Scale bar is 1 mm.
(C) Coronal section showing expression of DIO-hM4D(G_i_)-mCherry (red) terminals in the GPe and sites of electrolytic lesion. Scale bar is 1 mm.
(D) – **(E)** Representative single units in the GPe before, during, and after striatal infusion of saline (D) or CNO (E). *Insets*: Average waveform for representative single unit. Scale bar is 20 μs.

**(F) – (I)** Summary of GPe single-unit at baseline and post-infusion of saline (F, H; N = 2, n = 3, WSR: p = 0.5) or CNO (F, I; N = 2, n = 7, WSR: p = 0.9375) in the striatum. Firing rate is normalized to baseline activity. Vertical dashed lines represent infusion start and stop times.

**(J) – (K)** Representative single units in the GPe before, during, and after intraperitoneal (IP) injection of saline (j) or CNO (k). *Insets*: Average waveform for representative single unit. Scale bar is 20 μs.

**(L) – (O)** Summary of GPe single-unit activity at baseline and post-injection of saline (L, N; N = 3, n = 9, WSR: p = 0.0039) or CNO (M, O; N = 2, n = 8, WSR: p = 0.0078). Firing rate is normalized to baseline activity. Vertical dashed line represents time of IP injection. N = animals, n = single-units. Data shown as mean ± SEM.

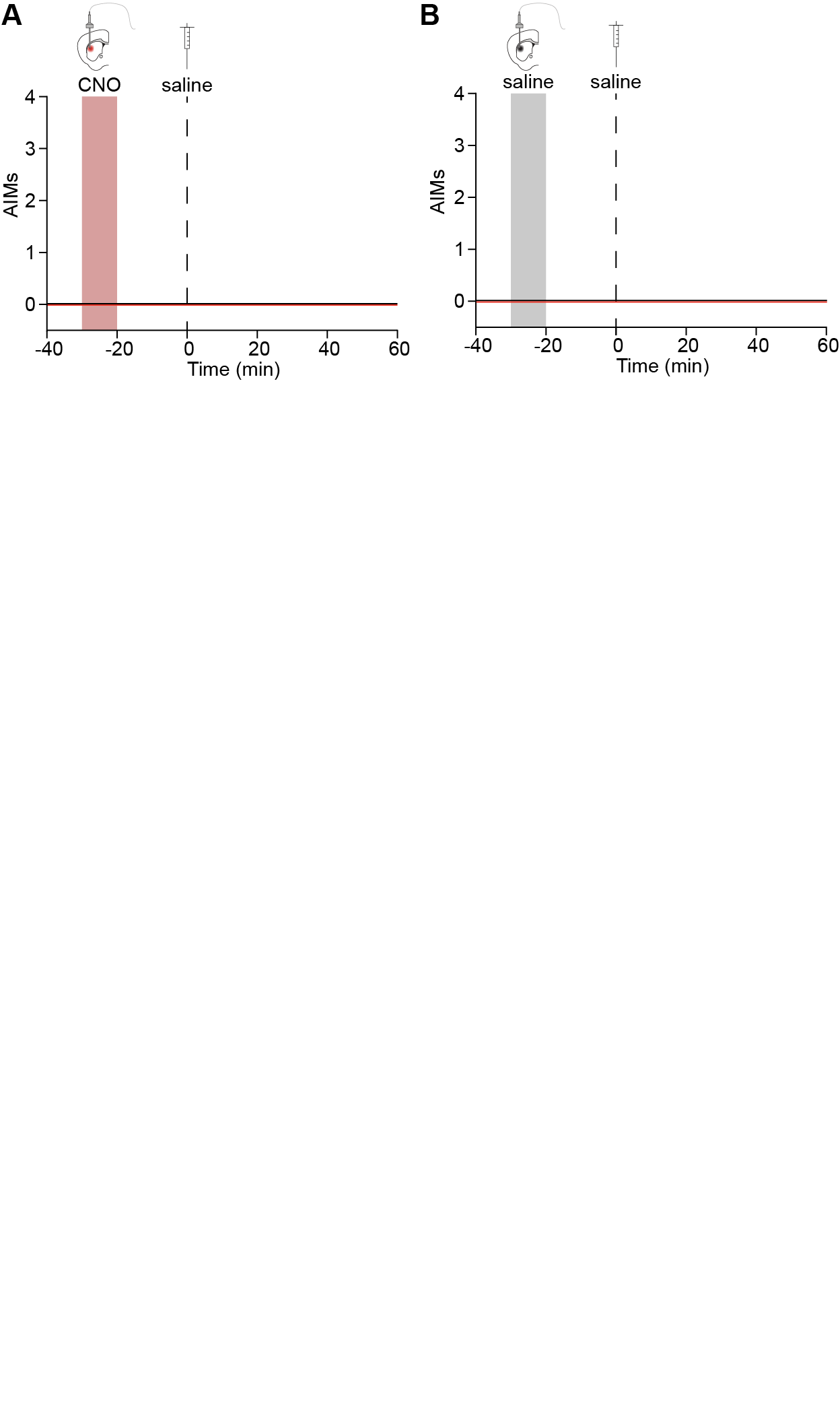

**Figure S7.** *Chemogenetic inhibition of striatal lateral connections originating from D2-MSNs is not sufficient to cause dyskinesia without levodopa, related to Figure 4.*

(A) – **(B)** No dyskinesia was observed following local striatal infusion of CNO (A) or saline (B) and IP injection of a low-dose of levodopa hM4D(G_i_): N = 14, mCherry: N = 12. N = animals. Data shown as mean ± SEM.

| **Key Experiments** | **Figure** | **Type of Comparison** | **Statistical Test** | **n (cells)** | **N (animals)** | **Comparison values (± SEM) & p-value** |
| --- | --- | --- | --- | --- | --- | --- |
| **Ctrl oIPSC amplitude, 1 mW (all groups)** | 1g | Between-group | KW | D2-D1: 41; D1-D1: 13; D1-D2: 20; D2-D2: 8 | D2-D1: 12; D1-D1: 4; D1-D2: 9; D2-D2: 5 | Χ^2^ = 19.25; p = 0.0002 |
| **Ctrl oIPSC amplitude (D2-D1 vs D1-D1)** | 1g | Between-group | Dunn’s | D2-D1: 41; D1-D1: 26 | D2-D1: 12; D1-D1: 5 | D2-D1 vs D1-D1, p = 0.0190 |
| **Ctrl oIPSC amplitude (D2-D1 vs D1-D2)** | 1g | Between-group | Dunn’s | D2-D1: 41; D1-D2: 20 | D2-D1: 12; D1-D2: 9 | D2-D1 vs D1-D2, p = 0.0012 |
| **Ctrl oIPSC amplitude (D2-D1 vs D2-D2)** | 1g | Between-group | Dunn’s | D2-D1: 41; D2-D2: 13 | D2-D1: 12; D2-D2: 6 | D2-D1 vs D2-D2, p = 0.0277 |
| **Ctrl oIPSC amplitude (D1-D1 vs D1-D2)** | 1g | Between-group | Dunn’s | D1-D1: 26; D1-D2: 20 | D1-D1: 5; D1-D2: 9 | D1-D1 vs D1-D2, p > 0.9999 |
| **Ctrl oIPSC amplitude (D1-D1 vs D2-D2)** | 1g | Between-group | Dunn’s | D1-D1: 26; D2-D2: 8 | D1-D1: 5; D2-D2: 5 | D1-D1- vs D2-D2, p > 0.9999 |
| **Ctrl oIPSC amplitude (D1-D2 vs D2-D2)** | 1g | Between-group | Dunn’s | D1-D2: 20; D2-D2: 13 | D1-D2: 9; D2-D2: 6 | D1-D2 vs D2-D2, p > 0.9999 |
| **MSN-MSN oIPSC amplitude after picrotoxin** | S1 | Within-cell | WSR | 16 | 13 | p < 0.0001 |
| **D1-D1 oIPSC amplitude, 1 mW (all groups)** | 2g | Between-group | KW | Ctrl: 26; Park: 28; LD: 24 | Ctrl: 5; Park: 7; LD: 8 | Χ^2^ = 8.512; p = 0.0142 |
| **D1-D1 oIPSC amplitude (Ctrl vs Park)** | 2g | Between-group | Dunn’s | Ctrl: 26; Park: 28 | Ctrl: 5; Park: 7 | Ctrl vs Park, p = 0.0452 |
| **D1-D1 oIPSC amplitude (Ctrl vs LD)** | 2g | Between-group | Dunn’s | Ctrl: 26; LD: 24 | Ctrl: 5; LD: 8 | Ctrl vs LD, p = 0.0271 |
| **D1-D1 oIPSC amplitude (Park vs LD** | 2g | Between-group | Dunn’s | Park: 28; LD: 24 | Park: 7; LD: 8 | Park vs LD, p > 0.9999 |
| **D2-D1 oIPSC amplitude, 1 mW (all groups)** | 2i | Between-group | KW | Ctrl: 41; Park: 24; LD: 32 | Ctrl: 12; Park: 6; LD: 8 | Χ^2^ = 14.68; p = 0.0006 |
| **D2-D1 oIPSC amplitude (Ctrl vs Park)** | 2i | Between-group | Dunn’s | Ctrl: 41; Park: 24 | Ctrl: 12; Park: 6 | Ctrl vs Park, p = 0.0288 |
| **D2-D1 oIPSC amplitude (Ctrl vs LD)** | 2i | Between-group | Dunn’s | Ctrl: 41; LD: 31 | Ctrl: 12; LD: 8 | Ctrl vs LD, p = 0.3685 |
| **D2-D1 oIPSC amplitude (Park vs LD)** | 2i | Between-group | Dunn’s | Park: 24; LD: 32 | Park: 6; LD: 8 | Park vs LD, p = 0.0004 |
| **D1-D2 oIPSC amplitude, 1mW (all groups)** | S3 | Between-group | KW | Ctrl: 20; Park: 30; LD: 16 | Ctrl: 9; Park: 4, LD: 9 | Χ^2^ = 0.8926; p = 0.64 |
| **D1-D2 oIPSC amplitude (Ctrl vs Park)** | S3 | Between-group | Dunn’s | Ctrl: 20; Park: 30 | Ctrl: 9; Park: 4 | Ctrl vs Park, p > 0.9999 |
| **D1-D2 oIPSC amplitude (Ctrl vs LD)** | S3 | Between-group | Dunn’s | Ctrl: 20; LD: 16 | Ctrl: 9; LD: 9 | Ctrl vs LD, p > 0.9999 |
| **D1-D2 oIPSC amplitude (Park vs LD)** | S3 | Between-group | Dunn’s | Park: 30; LD: 16 | Park: 4; LD: 9 | Park vs LD, p > 0.9999 |
| **Norm. D2-D1 oIPSC after quinpirole (all groups)** | 3d | Between-group | KW | Ctrl: 11; Park: 14; LD: 9 | Ctrl: 6; Park: 7; LD: 3 | Χ^2^ = 0.1004; p = 0.9510 |
| **Norm D2-D1 oIPSC after quinpirole (Ctrl vs Park)** | 3d | Between-group | Dunn’s | Ctrl: 11; Park: 14 | Ctrl: 6; Park: 7 | Ctrl vs Park, p > 0.9999 |
| **Norm D2-D1 oIPSC after quinpirole (Ctrl vs LD)** | 3d | Between-group | Dunn’s | Ctrl: 11; LD: 9 | Ctrl: 6; LD: 3 | Ctrl vs LD, p > 0.9999 |
| **Norm D2-D1 oIPSC after quinpirole (Park vs LD)** | 3d | Between-group | Dunn’s | Park: 14; LD: 9 | Park: 7; LD: 3 | Park vs LD, p > 0.9999 |
| **Ctrl D2-D1 oIPSC amplitude after quinpirole** | 3d | Within-Cell | WSR | 11 | 6 | p = 0.001 |
| **Park D2-D1 oIPSC amplitude after quinpirole** | 3d | Within-Cell | WSR | 14 | 7 | p = 0.0001 |
| **LD D2-D1 oIPSC amplitude after quinpirole** | 3d | Within-Cell | WSR | 9 | 3 | p = 0.0039 |
| **D2-D1 oIPSC Baseline vs Quinpirole vs Sulpiride** | S4b | Within-Cell | Friedman | 8 | 3 | Χ^2^ = 13; p = 0.0003 |
| **Norm D2-D1 oIPSC baseline vs quinprole** | S4b | Within-Cell | Dunn’s | 8 | 3 | Bsln vs Quin, p = 0.0014 |
| **Norm D2-D1 oIPSC baseline vs sulpiride** | S4b | Within-Cell | Dunn’s | 8 | 3 | Bsln vs Sulp, p = 0.9519 |
| **Norm D2-D1 oIPSC quinpirole vs sulpiride** | S4b | Within-Cell | Dunn’s | 8 | 3 | Quin vs Sulp, p = 0.0373 |
| **D2-D1 oIPSC amplitude (w/ hM4Di) after CNO** | 4c | Within-Cell | WSR | 11 | 4 | p = 0.010 |
| **AIMs: CNO infusion + IP levodopa (hM4Di vs mCherry)** | 4e | Between-group | 2-way RM ANOVA | NA | hM4Di: 14, mCherry: 12 | p < 0.001 |
| **AIMS: IP CNO + IP levodopa (hM4Di vs mCherry)** | 4i | Between-group | 2-way RM ANOVA | NA | hM4Di: 14, mCherry: 12 | p = 0.0059 |
| **D2-MSN RMP (bsln vs CNO)** | S5a | Within-Cell | WSR | 11 | 3 | p = 0.1309 |
| **D2-MSN rheobase (bsln vs CNO)** | S5b | Within-Cell | WSR | 11 | 3 | p = 0.3594 |
| **D2-MSN AP threshold (bsln vs CNO)** | S5c | Within-Cell | WSR | 11 | 3 | p = 0.3223 |
| **D2-MSN Maximal firing rate (bsln vs CNO)** | S5d | Within-Cell | WSR | 11 | 3 | p = 0.5566 |
| **D2-MSN Input resistance (bsln vs CNO)** | S5e | Within-Cell | WSR | 11 | 3 | p = 0.4131 |
| **DLS Infusion: Saline, GPe single-unit (pre- vs post-infusion)** | S6h | Within-Unit | WSR | 3 | 2 | p = 0.5 |
| **DLS Infusion: CNO, GPe single-unit (pre- vs post-infusion)** | S6i | Within-Unit | WSR | 7 | 2 | p = 0.9375 |
| **IP Injection: Saline, GPe single-unit (pre- vs post-injection)** | S6n | Within-Unit | WSR | 9 | 3 | p = 0.0039 |
| **IP Injection: CNO, GPe single-unit (pre- vs post-injection)** | S6o | Within-Unit | WSR | 8 | 2 | p = 0.0078 |

**Table S1.** *Statistical table.*

Each row in the table provides key information regarding a statistical comparison made in the manuscript, including the figure, statistical test, n/N, p-value, and planned sample size based on power calculations. Abbreviations: KW (Kruskal-Walis Test), Dunn’s (Dunn’s Multiple Comparisons Test), WSR (Wilcoxon Signed Rank), 2-way RM ANOVA (2-way repeated measures analysis of variance), Friedman (Friedman Test), MU (multi-unit). *p-value corrected for multiple comparisons.
